# Spatial Dynamics of the Developing Human Heart

**DOI:** 10.1101/2024.03.12.584577

**Authors:** Enikő Lázár, Raphaël Mauron, Žaneta Andrusivová, Julia Foyer, Mengxiao He, Ludvig Larsson, Nick Shakari, Sergio Marco Salas, Christophe Avenel, Sanem Sariyar, Jan N. Hansen, Marco Vicari, Paulo Czarnewski, Emelie Braun, Xiaofei Li, Olaf Bergmann, Christer Sylvén, Emma Lundberg, Sten Linnarsson, Mats Nilsson, Erik Sundström, Igor Adameyko, Joakim Lundeberg

## Abstract

Heart development relies on a topologically defined interplay between a diverse array of cardiac cells. We finely curated spatial and single-cell measurements with subcellular imaging-based transcriptomics validation to explore spatial dynamics during early human cardiogenesis. Analyzing almost 80,000 individual cells and 70,000 spatially barcoded tissue regions between the 5.5^th^ and 14^th^ postconceptional weeks, we identified 31 coarse- and 72 fine-grained cell states and mapped them to highly resolved cardiac cellular niches. We provide novel insight into the development of the cardiac pacemaker-conduction system, heart valves, and atrial septum, and decipher heterogeneity of the hitherto elusive cardiac fibroblast population. Furthermore, we describe the formation of cardiac autonomic innervation and present the first spatial account of chromaffin cells in the fetal human heart. We support independent exploration of our datasets by an open-access, spatially centric interactive viewer. In summary, our study delineates the cellular and molecular landscape of the developing heart’s architecture, offering links to genetic causes of heart disease.

## INTRODUCTION

The heart serves as the central organ of the cardiovascular system by generating a pressure gradient that enables blood circulation throughout the body. The sequential activation of two ventricles and two atria is orchestrated by the cardiac pacemaker-conduction system, while the tandem operation of two sets of cardiac valves blocks retrograde blood flow during the cardiac cycle. Heart rate and contraction force are also tightly regulated by the autonomic nervous system, adjusting cardiac output to the momentary needs of the body.

The heart forms its primary structures already within the 1^st^ trimester of prenatal development^1,2^. Cardiogenic mesoderm emerges during the 3^rd^ postconceptional week, which evolves into a linear heart tube by the end of the 5^th^ week. Through extensive spatial rearrangements and differentiation of diverse cell types, the cardiac tube transforms into the multilayered, four-chambered heart by the 8^th^ week, with maturation and growth continuing throughout fetal and postnatal development. Besides early cardiac precursors, other cells of extracardiac origin, including epicardial and neural crest cells, contribute to heart formation. Overall, early cardiogenesis follows a tightly regulated spatiotemporal progression, where disruptions can lead to congenital heart anomalies, highlighting the importance of understanding its governing molecular mechanisms^3,4^.

Anatomical changes throughout various stages of cardiogenesis have been studied in several species, including humans^5–10^. Over the past decade, the widespread adoption of single-cell RNA sequencing techniques yielded fresh insight into cardiac development on the cellular and molecular levels, describing transcriptomic profiles of cells within embryonic and fetal hearts, both in animal models^11–21^ and humans^22–29^. However, single-cell studies are inherently limited by their inability to retain the spatial context of isolated cells, which plays a pivotal role during morphogenesis. Recent advancements in spatial transcriptomics technologies allowed the detection of RNA expression within tissue sections, capturing previously inaccessible information on molecular arrangements in two dimensions. Combining these two technologies enables a more nuanced understanding of cell identities and interactions, factoring in their transcriptomic signature and their position within the tissue^30,31^. This approach has been successfully employed in a recent study investigating cardiogenesis in chicken^20^, the first published spatiotemporal atlas of human heart development^32^, and a recent report assessing cellular communities in the fetal human heart^33^.

Here we present a deep spatiotemporal map of the developing human heart during the 1st and early 2nd trimester. We analyzed 38 hearts between the 5.5^th^ and 14^th^ postconceptional weeks and assembled an extensive dataset of 69,114 spatially barcoded tissue spots and 76,801 individual cells, complemented by spatial detection of 150 selected transcripts by *in situ* sequencing (ISS). We discerned 23 spatial compartments with distinct transcriptional profiles within the cardiac tissue, uncovering molecular factors of regionality. Furthermore, we identified 11 primary cell types and 72 fine-grained cell states, which we then mapped to corresponding regions in cardiac tissue sections, enabling their highly refined, spatially-aware annotation. This allowed us to characterize distinct components of the cardiac pacemaker-conduction system, investigate their interactions with the emerging autonomic innervation, and describe a novel resident chromaffin cell population within the fetal heart. We also assessed position-related endothelial and mesenchymal cell heterogeneity in the developing cardiac valves and atrial septum and described an array of spatially defined cell states in the mesenchymal cell-fibroblast compartment. By investigating the co-occurrence of different cardiac cell states, we also delineated the architecture of prominent cardiac niches, enabling their targeted analysis. Furthermore, we provide an interactive viewer (https://hdcaheart.serve.scilifelab.se/web/index.html) to facilitate independent exploration of our datasets and analysis results.

Our work substantially broadens our understanding of early cardiogenesis, and presents a novel approach to cell atlasing, by deciphering cell identities and tissue dynamics with focus on the spatial context.

## RESULTS

### Spatial Profiling of the Developing Human Heart

First, to create a comprehensive account on spatial molecular patterns during early cardiogenesis, we performed 10× Genomics Visium spatial transcriptomics analysis on 16 hearts collected between the 6^th^ and 12^th^ postconceptional weeks, complemented by *in situ* sequencing of 150 selected transcripts. We compiled a dataset of 69,114 tissue spots from 38 heart sections, covering all major structural components of the developing organ (Fig. 1A). For further analysis of molecular determinants of regionality, we selected 17 sections encompassing at least three cardiac chambers, and performed unsupervised clustering of the corresponding 25,208 tissue spots, resulting in 23 spatial clusters (Fig. 1A-B, Suppl. Fig. 1A, Suppl. Table 1). Two clusters aligned with blood remnants in the atria and ventricles (B_A, B_V), and one with myeloid cell transcriptomic signature (MY) appeared scattered across the myocardium, however, the remaining 20 clusters mapped to distinct cardiac regions (Fig. 1C-D, Suppl. Fig. 1A-B).

**Figure 1.**
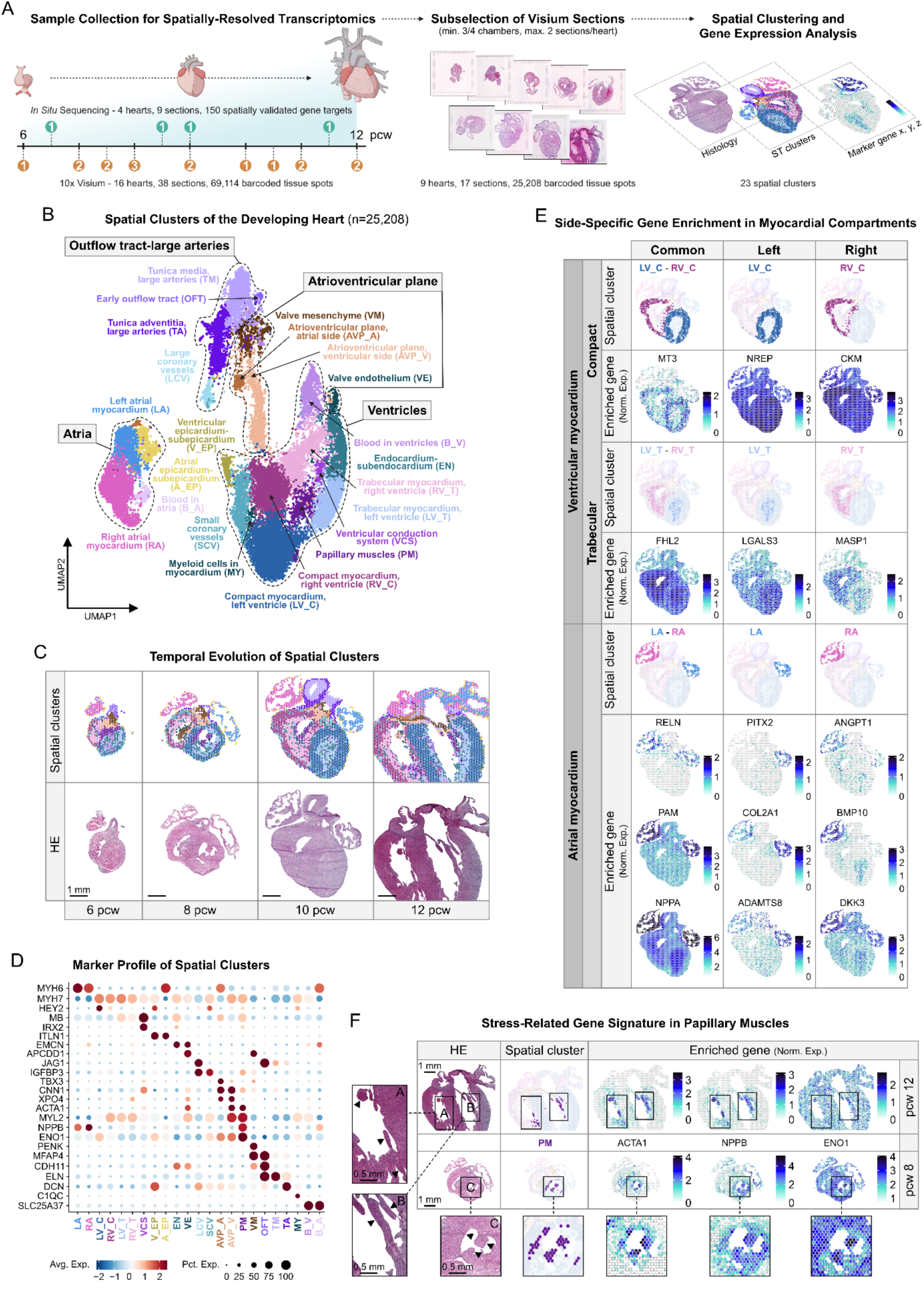
Spatial Profiling of the Developing Human Heart. **A**. Overview of spatially-resolved transcriptomic dataset generation, with donor numbers by postconceptional week (pcw) indicated in orange (Visium) and green (*in situ* sequencing) circles. **B.** 23 spatial clusters of the developing heart, corresponding to major cardiac structural components (dashed lines). **C.** Temporal evolution of spatial cluster distribution, presented in 6, 8, 10, and 12 pcw heart sections. Scale bar represents 1mm. **D.** Spatial feature plots displaying side-specific enrichment of selected DEGs of the left (LV_C) and right (RV_C) compact, left (LV_T) and right (RV_T) trabecular, and left (LA) and right (RA) atrial myocardial spatial clusters in a 10 pcw heart section. **E.** Spatial feature plots illustrating stress-related gene signature in papillary muscles (arrowheads in ROI A-C), featuring selected DEGs of the corresponding spatial cluster (PM). Scale bars represent 1 mm in the main, and 0.5 mm in the zoom-in panels. HE–hematoxylin-eosin.

Cardiac spatial clusters were broadly consistent across developmental stages. Still, we observed gradual disappearance of an early outflow tract-related cluster (OFT) and expansion of clusters representing the tunica media (TM) and adventitia (TA) of the developing large arteries (Suppl. Fig. 1C-D). The early OFT cluster’s transcriptomic profile substantially overlapped both with the TM and the valve mesenchyme-related VM clusters, likely due to their spatial proximity at early developmental stages. Notably, the VM-enriched gene *PENK*, a neural crest-derived mesenchymal cell marker described in mouse cardiac valves^34^, was not detected in the early OFT cluster, indicating a later contribution of this cell type to valve development (Fig. 1D, Suppl. Fig. 1A). Additional clusters represented the epicardial-subepicardial (A_EP, V_EP) and endocardial-subendocardial (EN) layers, small and large coronary vessels (SCV, LCV), and the developing valve endothelium (VE) (Fig. 1B-Dd, Suppl. Fig. 1A-B, E).

Our clustering approach delineated major myocardial compartments in a side-specific manner, facilitating the exploration of their transcriptomic differences (Fig. 1E, Suppl. Fig. 2A, E). Besides several common atrial markers, we observed selective enrichment of *PITX2*, *COL2A1*, and *ADAMTS8* in the left (LA), and *ANGPT1*, *BMP10*, and *DKK3* in the right atrial myocardial clusters (RA). The ventricle-associated spots were further divided into compact (LV_C, RV_C) and trabecular clusters (LV_T, RV_T), marked by the relative enrichment of *HEY2* and *MT3*, versus *MB* and *FHL2*, respectively. Furthermore, we detected side-specific enrichment *NREP* in the left, and *CKM* in the right compact layers, and of *LGALS3* and *IRX3* in the left, and *MASP1* and *PPP1R12B* in the right trabecular compartments. The transcription factor IRX3 is associated with ventricular conductive phenotype specification^35^, while galectin 3 (encoded by *LGALS3*) is established as a marker for adverse cardiac remodeling, primarily affecting the left ventricle^36^. A recent study proposed MASP1 as a candidate gene for ventricular conduction disorders^37^, and MYPT2 (encoded by *PPP1R12B*) is a known regulator of cardiomyocyte contraction force generation^38^. Side-specific enrichment of these genes might potentially contribute to the distinct electromechanical properties of the two ventricles.

Furthermore, we found a spatial cluster characterized by high expression of cardiomyocyte stress– and hypoxia response-related genes (*ACTA1*, *NPPB*, *ENO1*, *LDHA, MIF, FAM162A*)^39,40^ positioned around the papillary muscles (PM), reflecting mechanical strain on these structures (Fig. 1F, Suppl. Fig. 1A-B, E). Additionally, tissue spots with conduction system cell signatures localized to the ventricular subendocardium (VCS) and the atrial side of the atrioventricular plane (AVP_A), consistent with the position of bundle branches and atrioventricular nodal tissue, respectively (Fig. 1b-c, Suppl. Fig. 1a-b, e). Notably, we also observed a ventricular cluster of ambiguous identity, localized close to the atrioventricular plane (AVP_V), sharing markers with both AVP_A (*CNN1*, *XPO4*) and PM-MB (*ACTA1*, *MYL2*) clusters (Fig. 1B-C, Suppl. Fig. 1A-B, E).

We complemented our investigation with a spatially aware, neighborhood-based region identification approach^41^, which provided a good agreement with the spatial clusters representing larger myocardial and vessel compartments, but by design, did not distinguish fine or spatially dispersed structural components. At the same time, this analysis recognized a spatial domain corresponding to components of the intracardiac ganglionated plexi (ICGP), not recognized by our original clustering strategy (Extended Figure 1A-E). Additionally, we also performed non-negative matrix factorization (NMF) on the entire Visium dataset to decipher 20 spatial gene modules covarying in our samples, also included in our interactive viewer for independent assessment.

In summary, spatial profiling of the developing heart revealed previously unappreciated transcriptomic signatures in the papillary muscles and the atrioventricular region and provided insight into the molecular composition of other cardiac compartments.

### Cellular Landscape of Early Human Cardiogenesis

Next, we explored the cellular composition of the developing cardiac architecture through generating a single-cell RNA sequencing dataset from 15 hearts between the 5.5^th^ and 14^th^ postconceptional weeks, using the 10x Genomics Chromium platform. Unsupervised clustering of the merged 76,801 high-quality cardiac cells defined 31 coarse– and 72 fine-grained cell states with distinct transcriptomic profiles (Suppl. Table 2-6), creating a common framework for cell state identification throughout the analyzed period. We used these clusters to deconvolve the spatial transcriptomics dataset, mapping the predicted spatial distributions of cell states within the developing heart sections, and informing a spatially-aware annotation strategy (Fig. 2A).

**Figure 2.**
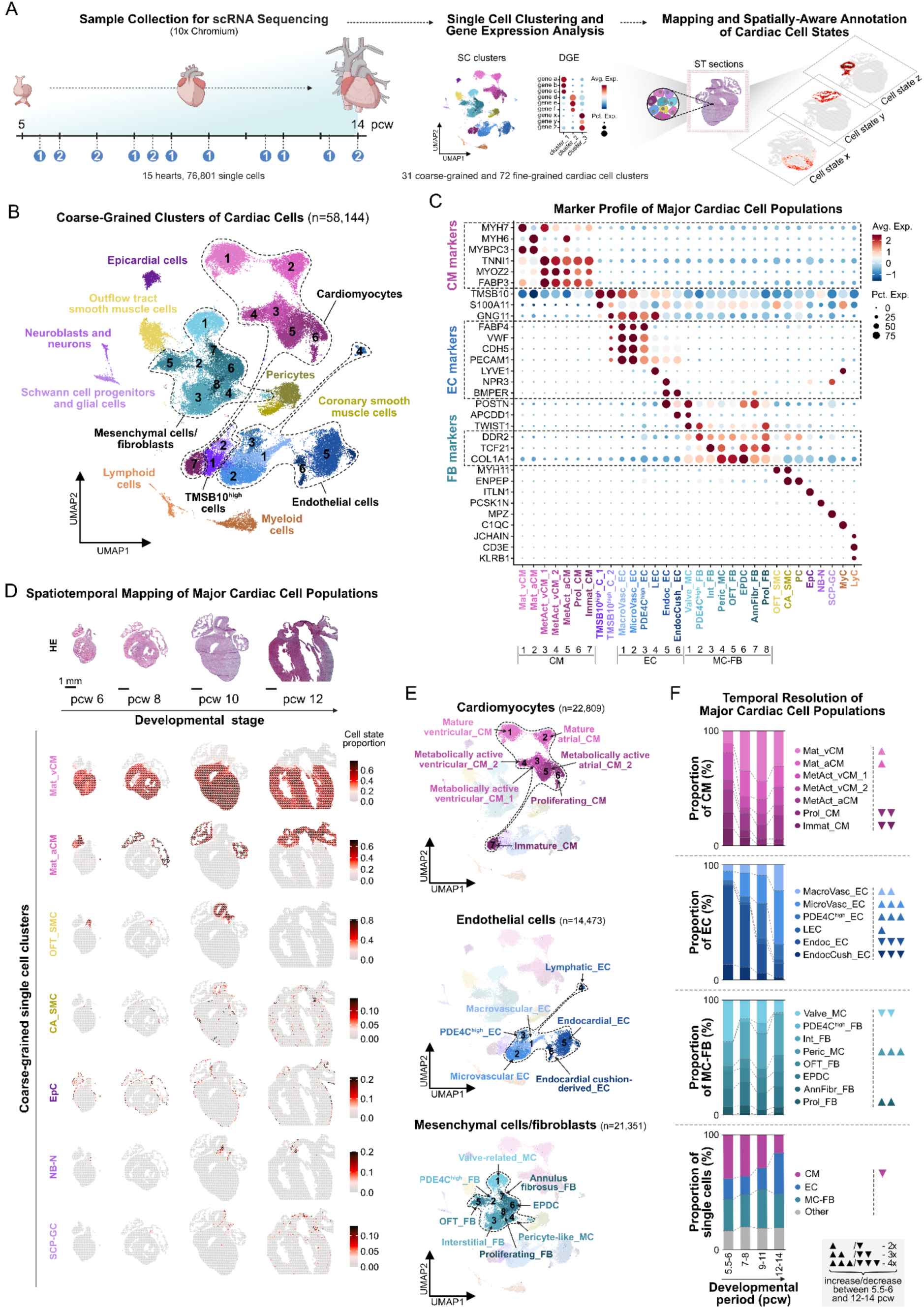
Cellular Landscape of Early Human Cardiogenesis. **A**. Overview of Chromium single cell dataset generation and spatially-informed annotation of cardiac cell states, with donor numbers by postconceptional week (pcw) shown in blue circles. **B.** UMAP representing 31 coarse-grained cardiac single-cell clusters, corresponding to 11 main cell types. **C.** Enrichment of canonical cell type markers across coarse-grained single-cell clusters. Dashed rectangles highlight consensus markers of cardiomyocytes (CM), endothelial cells (EC) and fibroblasts (FB). **D.** Spatiotemporal mapping of selected coarse-grained clusters in 6, 8, 10, and 12 pcw heart sections. Scale bar represents 1 mm, HE– hematoxylin-eosin. **E.** Spatially-aware annotation of coarse-grained cardiomyocyte, endothelial cell, and mesenchymal cell-fibroblast clusters. **F.** Temporal changes in coarse-grained cluster proportions in the cardiomyocyte, endothelial cell, and fibroblast-mesenchymal cell subsets across four developmental age groups (5.5-6, 7-8, 9-11, and 12-14 pcw). ▴-2x, ▴▴-3x, ▴▴▴-4x increase; ▾-2x, ▾▾-3x, ▾▾▾-4x decrease between the 5.5-6 and 12-14 pcw age groups.

Among the 31 coarse-grained clusters, expression of canonical cell type markers guided the identification of 11 main cell types, including predominant populations of cardiomyocytes, endothelial cells, and non-mural mesenchymal cells and fibroblast (Fig. 2B-C, Suppl. Table 2). We observed two distinct clusters expressing the smooth muscle cell marker *MYH11*, with one sharing transcriptional characteristics with a separate pericyte population (PC) (Fig. 2C, Suppl. Table 2, Ext. Fig. 2A-F, Ext. Text). Spatial mapping revealed distinct positions of these cell states in the outflow tract and great arteries (OFT_SMC) and large coronary arteries (CA_SMC), respectively (Fig. 2D). Additionally, we annotated epicardial cells (EpC), neuroblasts and neurons (NB-N), and Schwann cell progenitors and glial cells (SCP-GC), based on their marker expression and predicted positions on the epicardial surface, and around the great arteries and atrial walls, respectively (Fig. 2B-D, Suppl. Table 2). We also recognized two smaller populations of myeloid (MyC) and lymphoid cells (LyC), along with two clusters dominated by red blood cell transcriptomic signature, which we excluded from downstream analysis (Fig. 2B-C, Suppl. Table 2, Ext. Fig. 3A-E, Ext. Text). Furthermore, we also observed two clusters (TMSB10^high^_C_1-2) enriched in various thymosin transcripts, previously implicated in coronary vessel development^20,42^ (Fig. 2B-C, Suppl. Table 2).

The seven coarse-grained cardiomyocyte clusters displayed prominent differences in the expression of maturation, metabolic state, and cell cycle markers (Fig. 2C, E, Suppl. Fig. 2A), based on which we identified mature (Mat_vCM, Mat_aCM) and metabolically active (MetAct_aCM, MetAct_vCM_1-2) atrial and ventricular clusters, besides proliferating cardiomyocytes (Prol_CM). Additionally, we observed a population with lower cardiomyocyte-specific gene expression and a dynamic decrease in proportion (15.18% to 1.63%) over the investigated time frame, outlining immature cardiomyocytes (Immat_CM) (Fig. 2E-F, Suppl. Fig. 2A). Transcriptome-based identities of coarse-grained endothelial cell clusters were in agreement with their predicted localization in the tissue, outlining endocardial (Endoc_EC) and endocardial cushion-related cells (EndocCush_EC), macro-(MacroVasc_EC) and microvascular (MicroVasc_EC) endothelial cells of the coronary vasculature, and lymphatic endothelial cells (LEC) (Fig. 2C, E, Suppl. Fig. 2B). Specific markers for mesenchymal cell– and fibroblast subtypes in the fetal heart are currently lacking, thus positional cues are especially valuable to decipher cell identities. Using a spatially informed annotation strategy, we identified distinct populations of fibroblasts around the outflow tract and developing great arteries (OFT_FB), cardiac valve-related mesenchymal cells (Valve_MC), annulus fibrosus fibroblasts (AnnFibr_FB), interstitial fibroblasts (Int_FB) dispersed across the entire myocardium. We also observed a coarse-grained cluster representing epicardium-derived progenitor cells (EPDC) located in the subepicardial domain, with transcriptomic signatures consistent with an epicardial origin and ongoing epithelial-to-mesenchymal cell transition (EMT) (Fig. 2C, E, Suppl. Fig. 2B, Ext. Fig. 4A-D, Ext. Text). Furthermore, we recognized an additional mesenchymal cell population resembling cardiac pericyte transcriptomic profile (Peric_MC), and a cluster enriched in cell cycle genes (Prol_FB) (Fig. 2C, E, Suppl. Table 2).

Our clustering also highlighted an endothelial (PDE4C^high^_EC) and a fibroblast population (PDE4C^high^_FB) with overlapping transcriptomic profiles, enriched in several regulators of primary cilia formation and function (*TTC21A, ARL13B, TULP2*), and the cAMP-signaling-related genes *PDE4C* and *ATF3* (Fig. 2C, E, Suppl. Table 2). Ciliary cAMP signaling is involved in the differentiation of various cell types^43^, and a recent study revealed PDE4C regulating ciliary cAMP signaling in murine kidney cells^44^. By performing immunostaining for the broad cilia marker ARL13B, the basal body and centrosome marker PCNT, and PDE4C or ATF3, we observed ciliation, as well as PDE4C and ATF3 protein expression across the entire fetal heart (Suppl. Fig. 2C) and found basal bodies as prominent subcellular localization for both ATF3 and PDE4C. Thus, our data indicate a unique ciliated cell population, spread across the fetal heart, where cilium-related cAMP signaling may play an important role.

Temporal changes of cell state distributions in our dataset followed the main events of early cardiogenesis, including a shift towards mature cardiomyocyte profiles, pronounced expansion of vascular endothelial and mural cell populations associated with coronary vessel formation, a relative shrinkage of the epicardial, endocardial and cushion-related cellular compartments, and an expansion of the Schwann cell precursor-glial cell population in the developing innervation (Fig. 2F, Suppl. Fig. 2D). Interestingly, TMSB10^high^_C_1 and TMSB10^high^_C_2 clusters, also enriched in *S100A11* and *GNG11*, respectively, showed opposite temporal trends, presumably signaling a shift towards endothelial commitment in these populations (Suppl. Fig. 2D).

Differential gene expression analysis across time-resolved subpopulations of spatially annotated, coarse-grained single-cell states revealed relevant temporal transcriptional transitions within the endothelial and non-mural mesenchymal cell-fibroblast populations (Ext. Fig. 5A-C, Ext. Text), and also facilitated the assessment of relevant spatiotemporal patterns, such as presented in the area of the outflow tract and great vessels, and the cardiac valves (Ext. Fig. 6A-C, Ext. Text). Furthermore, enrichment analysis of heart disease-associated gene panels across the identified coarse– and fine-grained cell states broadly validated our cell type annotation, and highlighted the early and widespread expression and complex involvement of the confirmed genetic determinants in various forms of these pathologies (Ext. Fig. 7A-B).

Taken together, combined analysis of the single-cell and spatial datasets enabled refined identification of topologically distinct cell states and molecular arrangements, providing novel insight into cellular diversity in the developing heart.

### Molecular Map of the Cardiac Pacemaker-Conduction System Development

Contractile cardiomyocytes (CM) are the basic functional units of the myocardium; however, their coordinated activity relies on specialized cardiomyocytes of the cardiac pacemaker-conduction system (CPCS), featuring intrinsic automacy and efficient electrical impulse conduction. During development, these cells gradually acquire distinct electrophysiological properties from the surrounding working myocardium. Molecular characteristics of CPCS cells have been studied in several animal models^45–50^ and adult human hearts^51^, however, formation of the human CPCS remains uncharted.

To bridge this gap, we performed fine-grained clustering of the cardiomyocyte population and identified 19 distinct cell states, confirming the recently reported transcriptional diversity of these cells^52^. Besides a range of contractile cardiomyocytes of ventricular (vCM_1-6) and atrial origins (Left_aCM, Right_aCM, and Cond_aCM, representing smooth-walled conduit atrial regions), we also recognized cell states of lower maturation levels (Immat_CM_1-3), and of cell cycle signature (Prol_CM_1-2) (Fig. 3A-B, Suppl. Fig. 3A-B, Suppl. Table 3).

**Figure 3.**
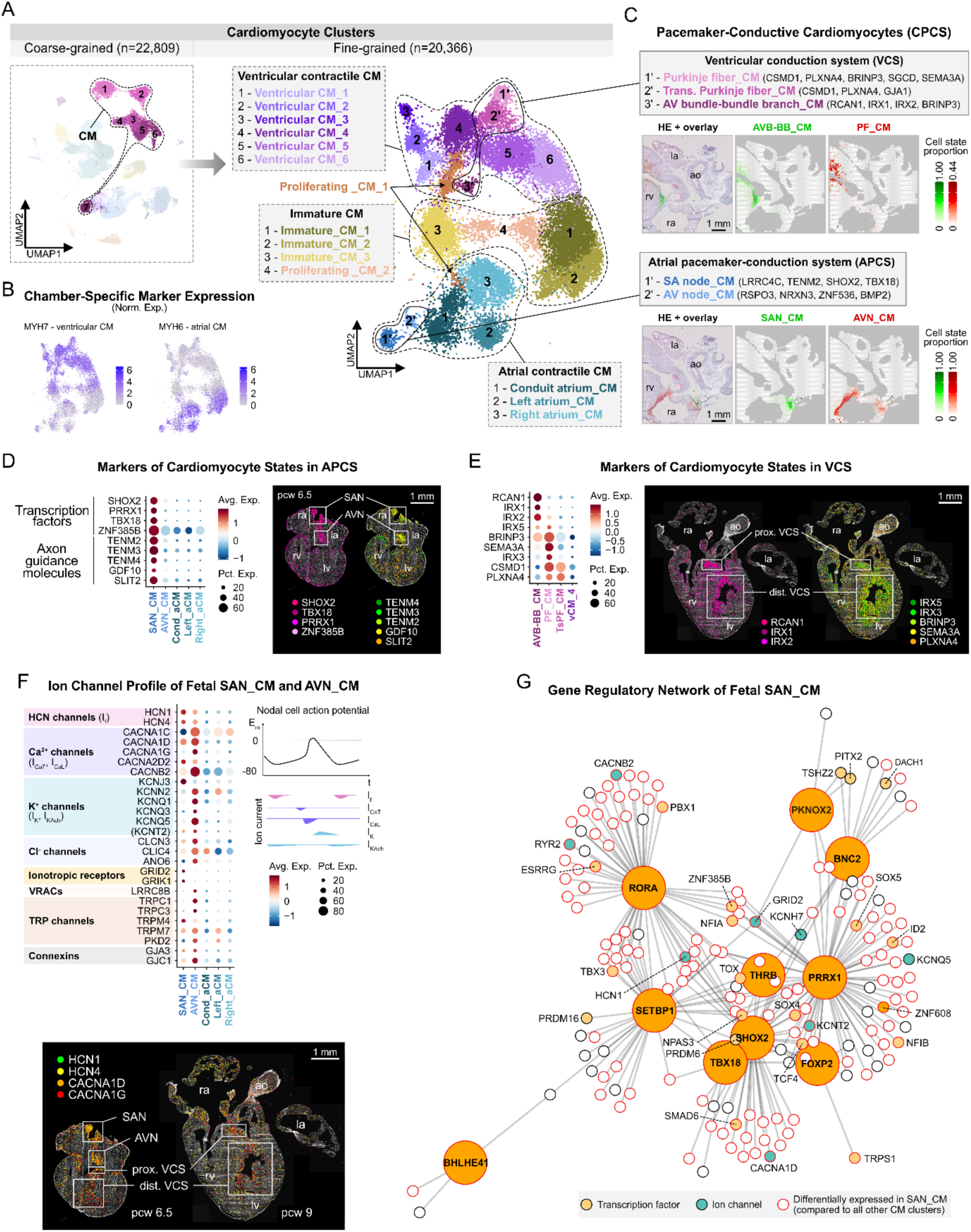
Molecular Map of Cardiac Pacemaker-Conduction System Development. **A**. UMAPs displaying 7 coarse-grained and 19 fine-grained cardiomyocyte clusters, representing ventricular and atrial contractile, immature, and pacemaker-conductive cell states. **B.** Feature plots of MYH7 and MYH6 outlining ventricular and atrial identities, respectively. **C.** Cardiomyocyte components of the cardiac pacemaker-conduction system (CPCS) with characteristic markers and predicted localization in an 11 pcw heart section. **D.** Dot plot representing selective enrichment of transcription factors and axon guidance molecules in SAN_CMs and AVN_CMs, compared to mature contractile atrial cardiomyocytes (left). Spatial enrichment of these genes in sinoatrial and atrioventricular nodes presented in a 6.5 pcw heart section by ISS (right). **E.** Dot plot illustrating the relative expression of CPCS marker genes between AVN-BB_CMs, PF_CMs, Ts-PF_CMs, and the contractile ventricular cardiomyocyte state vCM4 (left). Spatial expression patterns of these genes, detected by ISS in a 9 pcw heart section, corroborating their association with the proximal and distal ventricular conduction system (VCS) (right). **F.** Dot plot featuring relative expression of selected ion channels in SAN_CMs and AVN_CMs, compared to contractile atrial cardiomyocyte clusters (upper left). Illustration of ion currents of the nodal cell action (upper right). Spatial pattern of HCN– and T-type Ca^2+^ channel transcripts in 6.5 and 9 pcw heart sections, detected by ISS (lower). **G.** Gene regulatory network of SAN_CMs, including the top 10 DE transcription factors compared to all other cardiomyocyte cell states, and their associated target genes. In C-F panels: la–left atrium, ra–right atrium, rv–right ventricle, ao–aorta, SAN–sinoatrial node, AVN–atrioventricular node, VCS–ventricular conduction system, HE–hematoxylin-eosin; scale bars represent 1 mm.

Importantly, we observed clusters with robust expression of previously described markers of CPCS components (Fig. 3C, Suppl. Fig. 3A)^45–51^. Spatial mapping traced one such cluster to the upper wall of the right atrium, consistent with the sinoatrial node’s position (SAN_CM), while another appeared on the border zone between the atria and ventricles and in the atrial septum, delineating the developing atrioventricular node (AVN_CM). Two ventricular clusters featured markers of the atrioventricular bundle and bundle branches (AVB-BB_CM), and Purkinje fiber cardiomyocytes (PF_CM), and accordingly, were found in the ventricular subendocardium, stretching from the septum towards distal regions of the chambers (Fig. 3C). We observed an additional ventricular cluster with shared transcriptional characteristics and close spatial association to PF_CMs, representing transitional Purkinje fibers (Ts_PF) (Suppl. Fig. 3A, C).

CPCS cardiomyocytes displayed substantial overlaps in their transcriptomic profiles. Accordingly, SAN_CMs and AVN_CMs both expressed transcription factors specifically expressed (*SHOX2, TBX18*) or strongly enriched (*PRRX1, ZNF385B*) in pacemaker cardiomyocytes, and axon guidance molecules (*TENM2, TENM3, TENM4, GDF10, SLIT2*), outlining the nodes as essential targets of the developing cardiac innervation (Fig. 3D, Suppl. Fig. 3A). We also found the axon guidance molecule partner *LRRC4C*, previously not discussed in relation to the CPCS, specifically expressed in SAN_CMs (Suppl. Fig. 3A). Highly enriched genes in AVN_CMs reflected the node’s position in the atrioventricular plane (*BMP2*, *RSPO3*, *ADAMTS19)*, and neuronal characteristics of these cells (*NRXN3*, *ZNF536*) (Suppl. Fig. 3A). Among the ventricular CPCS cell states, PF_CMs showed expression of several characteristic markers (*CSMD1, BRINP3, SGCD, NTN1, SEMA3A*), partly shared with TsPF_CMs (Fig. 3E, Suppl. Fig. 3A). While recently described atrioventricular bundle markers *CNTN5* and *CRNDE* displayed broader distribution in our dataset, several genes enriched in AVB-BB_CMs, including *RCAN1* and *HS3ST3A1*, showed characteristic spatial enrichment in the upper edge of the ventricular septum, likely labeling the atrioventricular bundle (Fig. 3E, Suppl. Fig. 3D). Interestingly, the contractile vCM_1 cluster exhibited the highest *CNTN5* expression, beside other differentially expressed genes shared with CPCS (*TBX3, HS3ST3A1, BRINP3, TENM3*) and contractile ventricular cardiomyocyte states *(PRDM16, NLGN1, SORBS2, MYL2*) (Suppl. Fig. 3A, Suppl. Table 3). The gene enrichment profile (*XPO4*, *MYL2, TNFRSF19, SORBS2, ZFP36L1*) and predicted localization of vCM_1 cells also aligned with the elusive AVP_V spatial cluster, supporting a transitional ventricular cardiomyocytes identity of these cells, likely contributing to atrioventricular conductive tissue formation, as previously observed in the mouse heart^47^ (Suppl. Fig. 3B, E, Suppl. Table 1, 3).

Electrophysiological properties of CPCS components are determined by their ion channel repertoire^53^ (Fig. 3F, Suppl. Fig. 3F), which was recently explored in the adult human heart^51^. Similarly to their adult counterparts, developmental SAN_CMs and AVN_CMs featured ion channel profiles consistent with their nodal cell characteristics, including marked enrichment of hyperpolarization-activated cation channels *HCN1* and *HCN4*, and various L– and T-type Ca^2+^ channel genes (*CACNA1C, CACNA1D, CACNA1G, CACNA2D2, CACNB2*), responsible for the funny current (I_f_) and depolarization phase of the nodal action potential (I_Ca_), respectively. As opposed to adult cells, *CACNA1G* showed the highest enrichment in AVN_CMs, while SAN_CMs and ventricular CPCS cell states displayed similar, but somewhat lower expression of this gene. Repolarization in cardiomyocytes is primarily mediated by various potassium currents. Importantly, we found strong enrichment of the G-protein-coupled inwardly rectifying potassium (GIRK) channel subunit *KCNJ3* (but not of *KCNJ5*) in SAN_CMs, but not in PF_CMs, where the expression of this gene was described as a key distinctive feature from contractile ventricular cardiomyocytes in adult hearts^51^. On the other hand, PF_CMs showed high expression of *KCNJ2* and *KCNH7*, contributing to the inwardly rectifying (I_K1_) and fast delayed rectifier potassium currents (I_Kr_), respectively. AVN_CMs also featured strong expression of KCNQ1, previously associated with atrioventricular conduction block^54,55^, as well as several ion channels also present in contractile cardiomyocytes (*CACNA1C, KCNQ3, KCNQ5*), reflecting mixed electrophysiological properties of these cells. While *KCND2* (and the functionally related *KCND3*), previously described to be specifically enriched in the adult atrioventricular bundle, showed overall low expression in our dataset, a known interaction partner, *KCNIP4*, was strongly enriched in vCM_1 cells, providing further support to the contribution of this cell state to the developing atrioventricular conductive tissue. Gap junction-encoding gene enrichment showed a similar pattern as in adult hearts, with the low-conductance Cx45 (*GJC1*) marking SAN_CMs and AVN_CMs, and the high-conductance Cx40 (*GJA5*) being enriched in PF_CMs.

Beyond several chloride, volume-regulated anion (VRAC), and transient receptor potential (TRP) channels with yet unclear relevance in CPCS function, we also observed enrichment of several ionotropic glutamate receptor genes (*GRID2, GRIK1*) in SAN_CMs, in line with proposed glutamatergic signaling machinery in the adult node^51^, as well as of the GABA receptor ion channel *GABRB2* gene in Purkinje fibers, with the latter appearing specific to the developmental phase (Fig. 3F, Suppl. Fig. 3F).

To investigate the molecular drivers of pacemaker phenotype specification, we inferred a gene regulatory network of SAN_CMs (Fig. 3G). Key transcription factors (*SHOX2*, *PRDM6*, *THRB*, and *FOXP2*) and ion channel targets (*HCN1*, *CACNA1D*, *KCNT2*, *CACNB2*) exhibited substantial overlap between our results and a recent analysis of adult SAN cardiomyocytes^51^. Among other acknowledged pacemaker regulators enriched in SAN_CMs, such as *TBX18*, *BNC2*, and *TBX3*, the latter showed markedly higher expression in AVN_CM, AVB-BB_CM and vCM_1. TBX3 is known to suppress the atrial contractile gene program, thus its broader expression in CPCS components aligns with their progressive fate restriction, previously described in mouse^56,57^. Notably, the SAN_CM regulatory network also included transcription factors *BHLHE41* and *RORA*, recognized regulators of the mammalian circadian clock, which has been proposed to underlie time-of-day variation in arrhythmia susceptibility^58,59^.

### Early Formation of the Cardiac Autonomic Nervous System and Resident Chromaffin Cells

Heart function is tightly controlled by the autonomic nervous system, relaying sympathetic and parasympathetic signals to the cardiac tissue through the intracardiac ganglionated plexi (ICGP). Cells of the ICGP are derived from the cardiac neural crest, but the precise course of this process in humans is yet unknown^60^.

To explore the early development of local cardiac innervation, we reclustered the coarse-grained neuroblast-neuron (NB-N) and Schwann cell precursor-glial cell (SCP-GC) populations into ten fine-grained clusters (Fig. 4A, Suppl. Fig. 4A, Suppl. Table 4). Six clusters showed enrichment of Schwann cell precursor and glial cell transcription factors *SOX10* and *FOXD3*, and various levels of myelin-related genes *MPZ*, *PMP22*, and *MBP*, outlining five SCP states gradually obtaining more mature glial characteristics (SCP_1-5), as well as a population of myelinating Schwann cells (My_SC) (Fig. 4A-B, Ext. Fig. 8). Based on their enrichment in neuronal markers *PRPH* and *STMN2,* we also identified two clusters of autonomic neuroblasts and neurons (Aut_Neu_1-2), besides an intermediate bridge cell state from SCPs, characterized by high *ASCL1* expression (Fig. 4A-B, Ext. Fig. 8). Notably, we detected a cluster with strong enrichment of *CHGA*, *CHGB*, *PENK*, highlighting a local neuroendocrine chromaffin cell population (Chrom_C) in the cardiac tissue (Fig. 4A-B, Ext. Fig. 8). This is, to our knowledge, the first *in situ* observation of intracardiac chromaffin cells in the developing heart, which are not present in the mouse model, and thus appear to be human-specific. To elucidate transitions between the annotated cell states, we performed RNA velocity and pseudotime analysis on PCA embedding of the dataset, which confirmed a fork-like transition from early Schwann cell precursor states towards two parallel trajectories, consistent with the neuronal-chromaffin and glial differentiation paths (Fig. 4C-D, Ext. Fig. 8).

**Figure 4.**
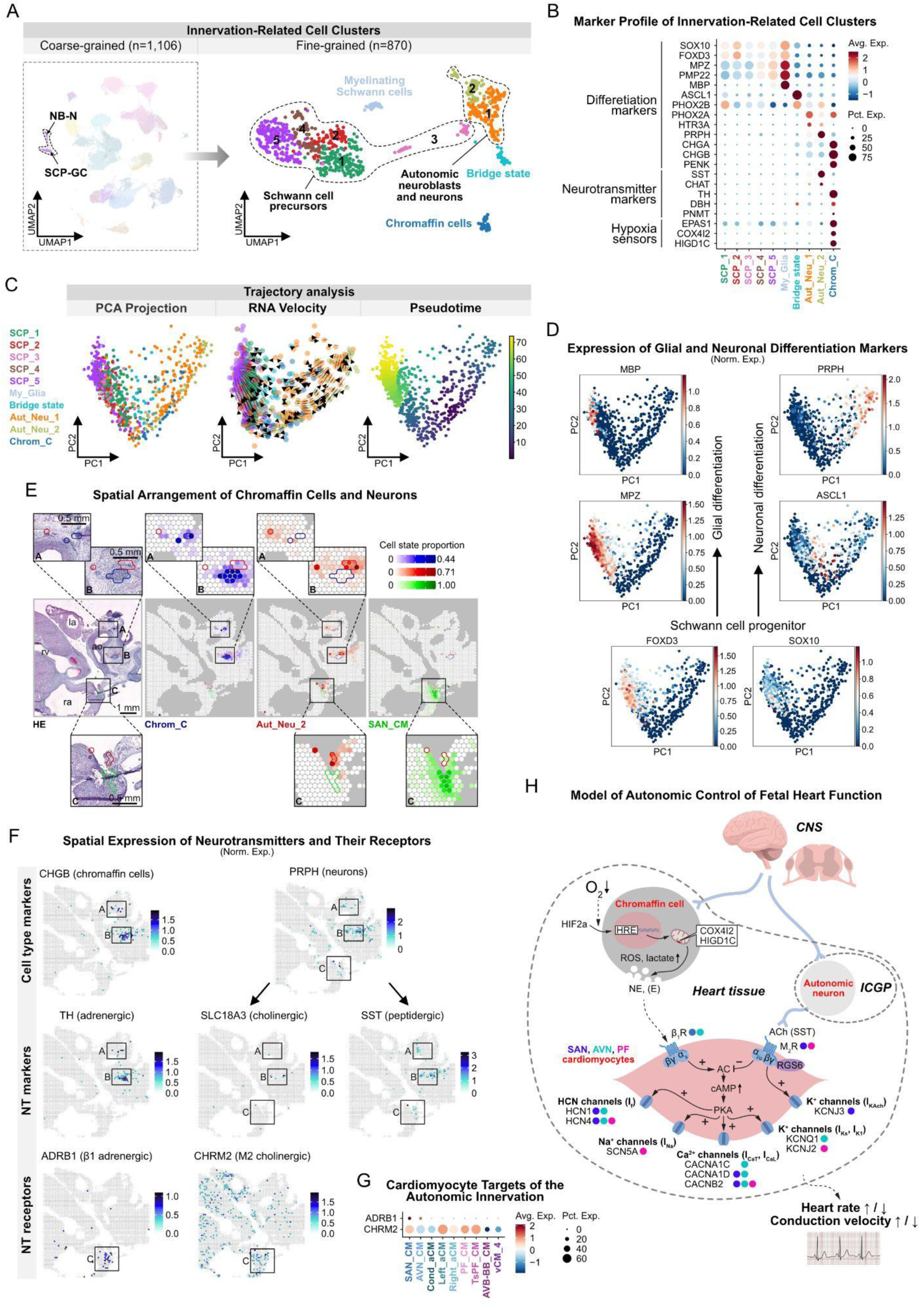
Early Formation of the Autonomic Cardiac Nervous System and Resident Chromaffin Cells. **A**. UMAPs showing the neuroblast-neuron (NB_N) and Schwann cell progenitor-glial cell (SCP-GC) coarse-grained, and 10 fine-grained clusters related to cardiac autonomic innervation. **B.** Dot plot illustrating the relative expression of glial, neuronal, and chromaffin cell differentiation markers, neurotransmitter-related genes, and components of the acute oxygen-sensing machinery, across all fine-grained innervation-related cell clusters. **C.** RNA velocity and pseudotime analysis revealing two parallel developmental trajectories among innervation-related cell clusters. **D.** Expression of glial and neuronal differentiation markers. **E.** Spatial mapping of Chrom_C (blue), Aut_Neu_2 (red) and SAN_CM (green) cell states in a 11 pcw heart section, outlining their neighboring compartments in the adventitia of the great vessels (ROI A-B), and an Aut_Neu_2 subset in proximity to the sinoatrial node (ROI C). Scale bars represent 1 mm in the main, and 0.5 mm in the zoom-in panels. **F.** Spatial feature plots of neurotransmitter metabolism-related genes and receptors in the same section, with consistent positions of ROI A-C. **G.** Dot plot presenting enrichment of β1 adrenergic and M2 cholinergic receptor transcripts in pacemaker-conduction system cardiomyocytes. **H.** Proposed model of autonomic control of fetal heart function. ICGP–intracardiac ganglionated plexi, SAN–sinoatrial node, AVN–atrioventricular node, PF–Purkinje fibers, CNS–central nervous system, ROS–reactive oxygen species. In panel D: la–left atrium, ra–right atrium, rv–right ventricle, ao–aorta, HE–hematoxylin-eosin.

In agreement with the predominant positions of ICGP, chromaffin cells and autonomic neurons were spatially mapped to the atrial wall and adventitia of the great arteries, where they appeared in closely associated, but not overlapping spatial domains, underscoring their common origin, but diverging functions (Fig. 4E, Suppl. Fig. 4B). Additionally, we observed Aut_Neu_2-dominated tissue segments in close proximity to SAN_CMs, highlighting a neuronal structure likely responsible for direct innervation of the nodal tissue (Fig. 4E). Most SCP cell states appeared more dispersed in the tissue, while the more mature SCP_5 cells and My_SCs concentrated around peripheral nerve structures (Suppl. Fig. 4C).

We identified somatostatin (*SST*) as the main neurotransmitter gene expressed in the Aut_Neu_1 and Aut_Neu_2 populations, besides the cholinergic neuron marker *CHAT* present in a smaller proportion of these cells. Meanwhile, the Chrom_Cs were strongly enriched in *TH*, and to a lower extent in *DBH* and *PNMT*, key genes of catecholamine synthesis (Fig. 4B, F). This suggests that neurons of the ICGP mostly function through peptidergic, and to a lesser degree, cholinergic transmission at this developmental stage, while the main source of norepinephrine (and to a lower extent, epinephrine) in the heart tissue is the local chromaffin cell population. Interestingly, we found negligible expression of somatostatin receptors in our datasets, meanwhile, β1 adrenergic receptor (*ADRB1*) was strongly enriched in the atrial, and M2 acetylcholine receptor (*CHRM2*) in almost all cardiomyocyte components of the pacemaker-conduction system (Fig. 4F-G). Distinct sources of adrenergic and cholinergic mediation of sinoatrial node function were reaffirmed by calculating a modified ligand-receptor score between the Chrom_C, Aut_Neu_2 and SAN_CM states, considering the coordinated expression levels of key enzymes (*TH* and *CHAT*) and receptors (*ADRB1* and *CHRM2*) between pairs of the analyzed cell states (Suppl. Fig. 4D). Additionally, genes involved in regulating neuronal migration, neurite growth, axon guidance, and synapse formation (such as teneurins, latrophilins, calsyntenins, neurexophilins, nectins, netrins, and components of the SNARE complex) showed specific enrichment in subsets of SAN_CMs and AVN_CMs, as well as in the Aut_Neu_1 and Aut_Neu_2 cell states (Suppl. Fig. 4E-F). Cell-cell communication analysis, based on ligands and receptors specifically enriched in the SAN_CM and Aut_Neu_2 cell states, highlighted the *PTPRS-NTRK3* interaction described in organizing excitatory synapses^61^, and the *TENM2-ADGRL1* pair known to induce axonal attraction on growth cones in the central nervous system^62^ (Suppl. Fig. 4G). These putative molecular interactions might be relevant for establishing precise connections between developing cardiac innervation and pacemaker cells.

An essential function of chromaffin cells is mediating a physiological response to hypoxia through increased catecholamine release. Accordingly, Chrom_Cs showed robust expression of molecular sensors for local oxygen tension, including *EPAS1* (encoding HIF2*α*), *COX4I2*, and *HIGD1C*, a mitochondrial electron transport chain component recently described in the oxygen-sensing machinery of the carotid body^63^ (Fig. 4B).

Based on this insight, we propose a cellular model of the functional interplay between the developing autonomic innervation, local chromaffin cells, and cardiomyocyte components of the early fetal human heart, where local neurons convey parasympathetic signals, and sympathetic modulation is largely mediated by a resident chromaffin cell population through a paracrine mechanism, in response to local tissue hypoxia. Resulting changes in heart function are likely conveyed through the activation of various ion channels in CPCS cardiomyocytes, adapting heart rate and conduction velocity to the environment of the developing fetus (Fig. 4H). An intracardiac population of chromaffin cells might also explain the origin and development of cardiac pheochromocytoma, a rare human condition when chromaffin cell tumors form in the cardiac region^64^.

### Position-Dependent Endothelial Cell Diversity in Endocardial Cushion-Derived Structures

Cardiac endothelial cells (EC) can be divided into vascular and endocardial subsets. With the swift growth of coronary vessels during the 1^st^ trimester, the vascular endothelium rapidly expands and diversifies. Meanwhile, the endocardium gives rise to endocardial cushions, contributing to the subsequent formation of cardiac valves, atrial septum and the upper membranous septum of the ventricles. The major cardiac endothelial populations were recently characterized in the fetal human heart^28^, however, we still lack a detailed, transcriptome-wide description of the scarce endocardial cushion-related endothelial cells and their various derivatives^23,65^.

In the merged population of coarse-grained endothelial cells and TMSB10^high^_C_2 clusters featuring endothelial transcriptional characteristics, we defined 18 fine-grained cell states, representing the endocardium (EndocEC_1-4), endothelium of large arteries and veins (Atr_EC_1 and Ven_EC), and consecutive segments of the coronary vasculature, including arterial (Art_EC_2), arteriolar (Arteriol_EC), capillary (Cap_EC_1-2) and venular (Venul_EC) endothelial cells. We also recognized separate populations of proliferating (Prol_EC) and lymphatic endothelial cells (LEC^fg^), as well as distinct populations of thymosin-(TMSB10^high^_C_2^fg^) and PDE4C-enriched cells (PDE4C^high^_EC) (Fig. 5A, Suppl. Fig. 5A-C, Suppl. Table 5).

**Figure 5.**
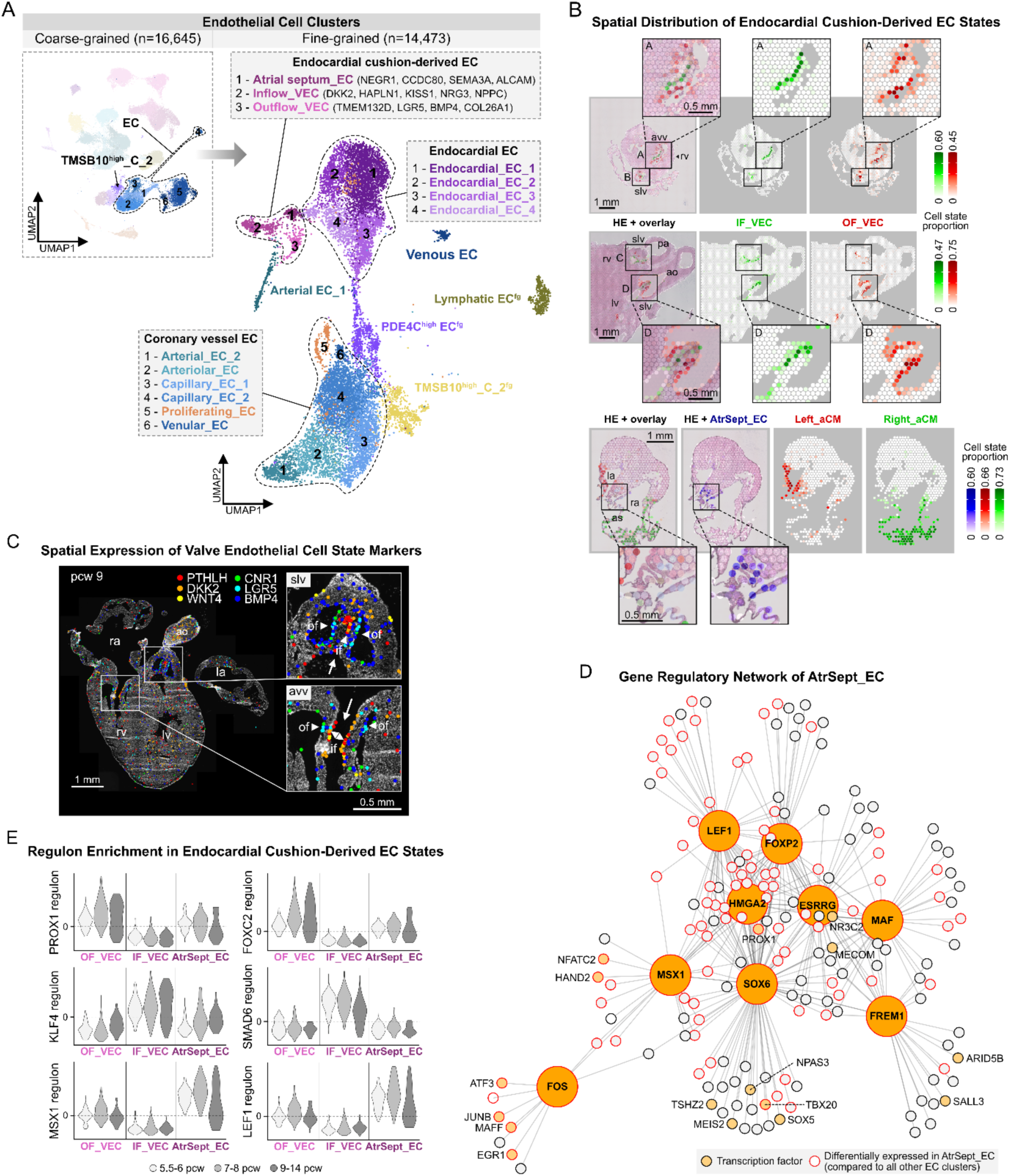
Position-Dependent Endothelial Cell Diversity in Endocardial Cushion-Derived Structures. **A**. UMAPs displaying 6 coarse-grained endothelial cell (EC)– and the TMSB10^high^_C_2 clusters, divided into 18 fine-grained cell states, representing endocardial, endocardial cushion-derived, and coronary vessel endothelial cells, along with additional minor populations. **B.** Spatial mapping tracing IF_VECs (green) and OF_VECs (red) to opposite sides of the atrioventricular (ROI A) and semilunar valves (ROI B-D) in 8 (upper) and 12 pcw (middle) heart sections, and AtrSept_ECs (blue) to the atrial septum between Right_aCM-(green) and Left_aCM (red)-dominated areas, in 8 pcw heart section (lower). Scale bars represent 1 mm in the main, and 0.5 mm in the zoom-in panels. **C.** Side-specific spatial enrichment of selected IF_VEC and OF_VEC markers in atrioventricular and semilunar valves in a 9 pcw heart section, detected by ISS. White arrows mark the direction of blood flow. Scale bar represents 1 mm. **D.** Gene regulatory network of AtrSept_EC, including enriched transcription factors compared to IF_VECs and OF_VECs, and their associated target genes. **E.** Violin plots illustrating regulon enrichment of selected transcription factors in AtrSept_ECs, IF_VECs, and OF_VECs across three age groups (5.5-6, 7-8, 9-14 pcw). In panels B-C: slv–semilunar valve, avv–atrioventricular valve, rv–right ventricle, lv– left ventricle, ao–aorta, pa–pulmonary artery, as–atrial septum, if–inflow side of valve, of–outflow side of valve, HE–hematoxylin-eosin.

Three additional clusters appeared closely related to endocardial cell states but were also enriched in key regulators of endothelial-to-mesenchymal transition (EndMT) (*NFATC1*, *APCDD1*, *TWIST1)*, central in endocardial cushion formation (Suppl. Fig. 5B, Suppl. Table 5). Accordingly, spatial mapping traced two of these cell states to opposite sides of the developing heart valves, outlining inflow (IF_VEC) and outflow (OF_VEC) valve endothelial cell populations, consistently in the atrioventricular and semilunar valves (Fig. 5B). This pattern was already apparent in primitive endocardial cushions from as early on as 6 postconceptional weeks, suggesting early specification of these populations (Suppl. Fig. 5D). Meanwhile, the third cluster showed highly distinct localization in the atrial septum (AtrSept_EC), illuminating a yet undescribed cardiac endothelial cell state (Fig. 5B, Suppl. Fig. 5E). Beyond several common markers, these three populations featured largely distinct transcriptional profiles (Suppl. Fig. 5A, F-G). Highly enriched genes of IF_VECs included several components of the WNT signaling axis (*WNT2*, *WNT4*, *WNT9B*, *DKK2*, *PTHLH*), with many of them displaying spatial enrichment in the coaptation zone of the valves (Fig. 5C). OF_VECs, on the other hand, displayed high expression of BMP ligands (*BMP4*, *BMP6*), WNT signaling modulators (*LGR5*), as well as the endocannabinoid receptor-encoding *CNR1*, showing spatial enrichment on the fibrosa side of the valves (Fig. 5C). AtrSept_ECs shared some highly enriched genes (*LRRC4C*, *LSAMP*, *OPCML)* with the Endoc_EC_4 cell state, outlining the smooth-walled atrium, besides more specific markers (*NEGR1*, *ALCAM*, *SEMA3A*, *CCDC80, MSX1*) (Suppl. Fig. 5A, E, Suppl. Table 5).

Gene regulatory network and regulon enrichment analysis gave further insight into the molecular differences between the three endocardial cushion-related cell states (Fig. 5D-E). We identified *MSX1* and *LEF1*, known mediators of EndMT during endocardial cushion formation, as the most highly enriched transcription factors in AtrSept_ECs. Besides the cardiac valve mesenchyme, *LEF1* is also expressed in the mesenchymal cap of the developing atrial septum in mice^66^, and a recent report described atrial septal defect in a patient carrying a heterozygous missense variant of this gene^67^. *MSX1* was also detected in the atrial septum of the mouse heart^68^, and several recent animal and human studies proposed a potential association between atrial septal defects and genetic variants of the *MSX1* locus^68–70^. In OF_VECs, we found enrichment of *PROX1* and *FOXC2* transcription factors, known to act in concert in response to oscillatory shear stress, maintaining extracellular matrix structure and preventing myxomatous degeneration of cardiac valves^71^. Several transcription factors enriched in IF_VECs have an established role in the modulation of WNT signaling, including *KLF4* and *SMAD6*, which are known transducers of laminar shear stress in endothelial cells^72,73^. Importantly, we observed pronounced differences in the regulon enrichment of these transcription factors between IF_VECs and OF_VECs from as early on as 5.5-6 postconceptional weeks, suggesting that hemodynamic forces, already in the embryonic period, are core factors in valve endothelial cell diversification.

### Spatial Decomposition of Fibroblast and Mesenchymal Cell Heterogeneity in the Developing Heart

Fibroblasts (FB) and mesenchymal cells (MC) constitute an ambiguous subset of cardiac cells during development, largely due to their pronounced transcriptional heterogeneity and a lack of well-established molecular markers. Spatial factors, reflecting the positions of their progenitors and niche-related environmental signals, play a central role in defining cellular identities in this population^74^.

To explore this topological diversity, we spatially mapped 18 fine-grained fibroblast and mesenchymal cell clusters distinguished in our dataset (Fig. 6A, Suppl. Fig. 6A-B, Suppl. Table 6), tracing most to characteristic tissue locations. Accordingly, we recognized two adventitial fibroblast states (Adv_FB_1-2), with Adv_FB_1 outlining the wall of the outflow tract and great arteries, and Adv_FB_2 also appearing around large coronary arteries (Fig. 6B). An additional mesenchymal cell population, resembling the pericyte transcriptional signature (Peric_MC^fg^), was traced to the innermost layer of the ventricular myocardium, as well as to the atrial walls, to positions complementary to the genuine pericyte population (Fig. 6B, Suppl. Fig. 6A-B). This localization is suggestive of a potential endocardial origin, which has recently been proposed for a pericyte-like cardiac mesenchymal cell state in mice^75^.

**Figure 6.**
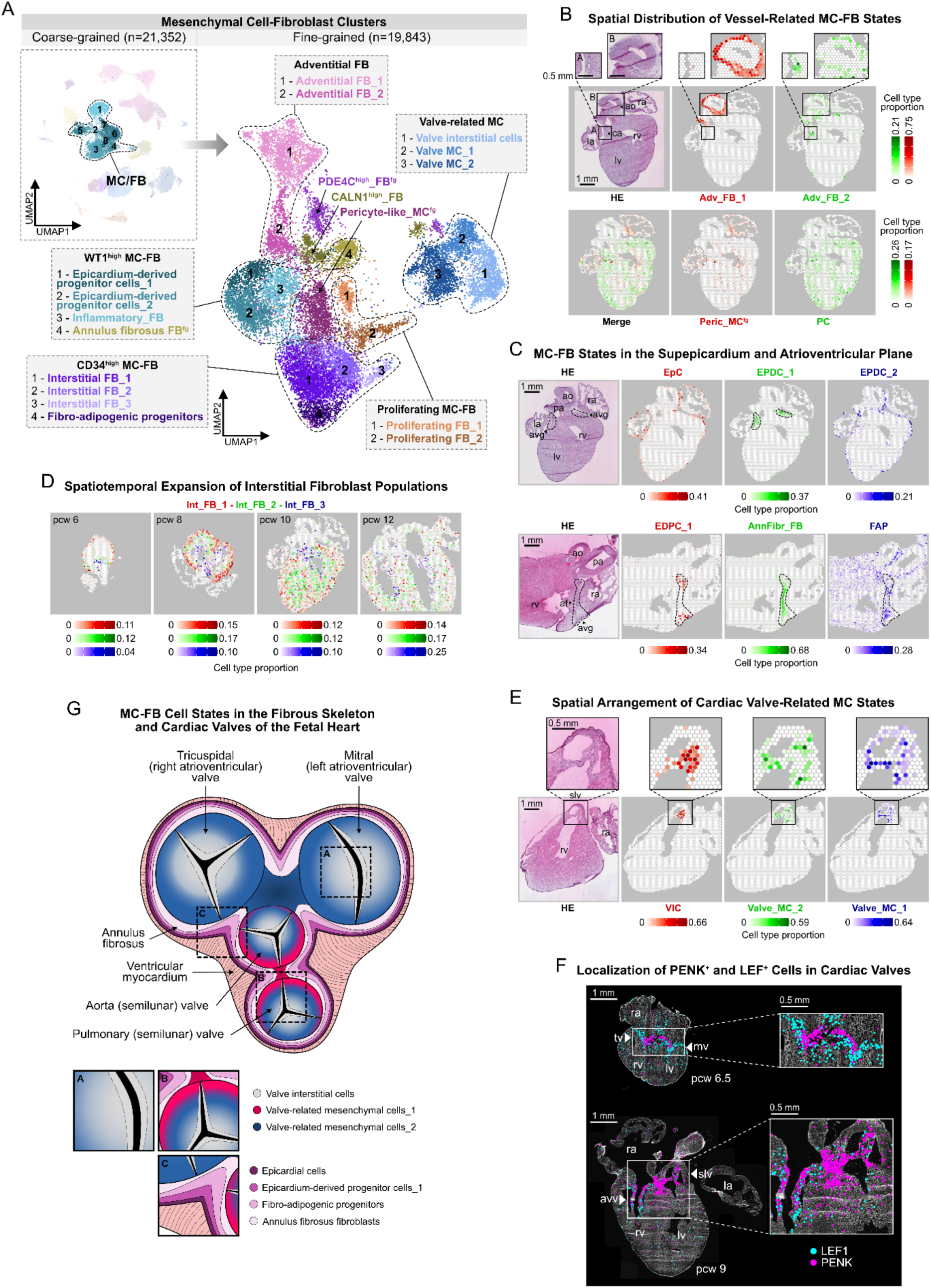
Spatial Decomposition of Fibroblast and Mesenchymal Cell Heterogeneity in the Developing Heart. **A**. UMAPs displaying 8 coarse-grained and 18 fine-grained fibroblast (FB) and mesenchymal cells (MC) clusters, representing WT1– and CD34-enriched, adventitial, valve-related and proliferating subsets, along with additional minor populations. **B.** Spatial mapping, tracing Adv_FB_1 (red) and Adv_FB_2 (green) cells to the great arteries (ROI B) and coronary arteries (ROI A), respectively (upper), and Peric_MC^fg^ (red) and PC (green) cells to complementary positions in the atrial and ventricular myocardium, in a 10 pcw heart section. **C.** Spatial mapping, tracing EpC (red), EPCD_1 (green), and EPCD_2 (blue) cells to the heart surface in a 10 pcw (upper), and EPCD_1 (red), AnnFibr_FB (green) and FAP (blue) cells to the atrioventricular plane and groove (dashed lines) in an 11 pcw heart section. **D.** Spatial mapping of Int_FB_1 (red), Int_FB_2 (green) and Int_FB_3 (blue) cells, illustrating their gradual spatial expansion and layer-specific positions in 6, 8, 10, and 12 pcw heart sections. **E.** Spatial mapping of VIC (red), Valve_MC_2 (green) and Valve_MC_1 (blue) cells in an 11 pcw heart section, displaying their enrichment in and around the semilunar valve cusps. **F.** ISS detection of *PENK* and *LEF1* in cardiac valves in 6.5 and 9 pcw heart sections. **G.** Spatial model of fibroblast and mesenchymal cell state arrangements in the fetal heart’s fibrous skeleton and cardiac valves. In panel B-C, E-F: rv–right ventricle, lv–left ventricle, ra–right atrium, la–left atrium, ao–aorta, ca–coronary artery, pa–pulmonary artery, avg– atrioventricular groove, af–annulus fibrosus, mv–mitral valve, tv–tricuspid valve, HE–hematoxylin-eosin; scale bars represent 1 mm in the main, and 0.5 mm in the zoom-in panels.

Furthermore, we identified two populations representing early forms of epicardium-derived progenitor cells (EDPC), based on their predicted positions in the subepicardium at the atrioventricular groove (EPDC_1) and the heart surface (EPDC_2), and combined enrichment of the epicardial marker *WT1* and EPDC marker *TCF21* (Fig. 6C, Suppl. Fig. 6B). A third *WT1*-enriched cell state, also expressing high levels of *CACNA2D3* and *BRINP3*, appeared in a distinct layer between the atria and ventricles, outlining the fibrous cardiac skeleton (AnnFibr_FB), beside a fourth population marked by inflammatory gene expression (Inf_FB) (Fig. 6C, Suppl. Fig. 6A). We also found various cell states consistent with an interstitial fibroblast phenotype, characterized by high *CD34* expression and gradual temporal expansion from the epicardium to the outer (Int_FB_1) and inner (Int_FB_2) layers of the ventricular wall, or located in the subendocardium and the atrioventricular region (Int_FB_3) (Fig. 6D, Suppl. Fig. 6B). Int_FB_1 and Int_FB_2 exhibited the highest *TCF21* expression in our dataset, indicating advanced transition from an epicardial progenitor state towards fibroblast identity (Suppl. Fig. 6B). Notably, we also recognized a distinct cell state with robust *CD34* expression, dynamic temporal expansion, and a transcriptomic profile resembling fibro-adipogenic progenitors (FAP; *CLEC3B, ROBO2, SEMA3C, ADAMTSL1, CCN3, SOX9, DLK1*) (Suppl. Fig. 6A-C). Many of these markers appeared closely associated in a gene regulatory network defined by three transcription factors enriched in this population: while *ZBTB16* promotes white and brown adipogenesis^76^, *KLF2* and *GLIS3* block further adipogenic differentiation of preadipocytes^77,78^, consistent with the progenitor characteristics of this cell state (Suppl. Fig. 6D). Spatial mapping located this population near EPDC_1 cells in the atrioventricular groove, reflecting the predominant localization of the primordial epicardial adipose tissue (EAT), as well as a presumed epicardial origin of these cells (Fig. 6C). FAPs have been proposed as potential sources of fatty-fibrous tissue deposits, a histological hallmark in arrhythmogenic right ventricular cardiomyopathy (ARVC)^79^. Importantly, we found two ARVC-associated desmosome-encoding genes (*PKP4*, *DSC2*) enriched in the FAP cell state, drawing a potential connection towards the pathogenesis of the disease^80^ (Suppl. Table 6).

Our clustering also identified three distinct mesenchymal cell states in the developing endocardial cushions and cardiac valves (Fig. 6E, Suppl. Fig. 6E-F). Valve interstitial cells (VIC; *APCDD1*, *LEF1*, *TMEM132C, ADAMTS19*) localized to the free segments of the valves, while the other two populations (Valve_MC_1-2) stretched from the intervalvular fibrous region towards the valve roots, with Valve_MC_1 (*FGF14*, *HDAC9*, *PLCXD3*) being spatially enriched around the semilunar valves. Importantly, Valve_MC_2 featured a specific and robust expression of *PENK*, a previously described marker of mesenchymal neural crest derivatives in mouse hearts^34^ (Suppl. Fig. 6A-B, E). Importantly, we observed strong spatial *PENK* signal in and around the developing semilunar and atrioventricular valves, supporting the long-debated contribution of neural crest cells to both valve structures in humans. In fact, we observed *PENK* expression in the septal leaflets of atrioventricular valves already in the 6.5^th^ postconceptional week, which appeared more spread out towards the free wall leaflets by the 9^th^ week, along with semilunar valve-related *PENK* signal present in both sampled aortic valve cusps (Fig. 6F). Taken together, our results imply a more substantial contribution of neural crest-derived mesenchyme to human cardiac valves than previously assumed. Based on these results, we assembled a spatial model of mesenchymal and fibroblast-like cell states contributing to the fibrous skeleton and embedded valve structures in the fetal heart, representing epicardial, endocardial and neural crest derivatives (Fig. 6G).

Beyond clusters with characteristic spatial distributions, we also identified a cell cycle marker-(Prol_FB) and a PDE4C-enriched cell state (PDE4C^high^_FB^fg^), beside a fibroblast population (CALN1^high^_FB) featuring high expression of characteristic nodal cell genes (*CALN1*, *SHOX2*, *CNTN5*), as well as the angiotensinogen-encoding *AGT*, a central component of local renin-angiotensin circuits in nodal tissue (Suppl. Fig. 6A, Suppl. Table 6). The latter observation highlights this population as potential developmental equivalents of sinoatrial and atrioventricular node-resident fibroblasts.

### Spatially Informed Analysis of Developmental Cardiac Niches

During cardiogenesis, rapid formation and rearrangements of cellular microcompartments play fundamental roles in conveying external signals and cues to cardiac cells, regulating their behavior in the evolving organ. Hence, our objective was to utilize our in-depth spatial analysis to outline the dominant cell types in major structural compartments and more refined niches in the developing heart, expecting that their focused interrogation would aid in understanding the cellular and molecular mechanisms that govern both normal and pathological heart formation. Accordingly, we assessed the overlap of predicted spatial distributions between the 72 fine-grained single-cell states in our dataset and used in-pair correlation-based co-detection scores to visualize their topological relations (Fig. 7A, Suppl. Table 7). With this strategy, we were able to decipher the cellular composition of larger and smaller tissue compartments consistently across three developmental age groups, confirming the robustness of our approach (Suppl. Fig. 7A).

**Fig. 7.**
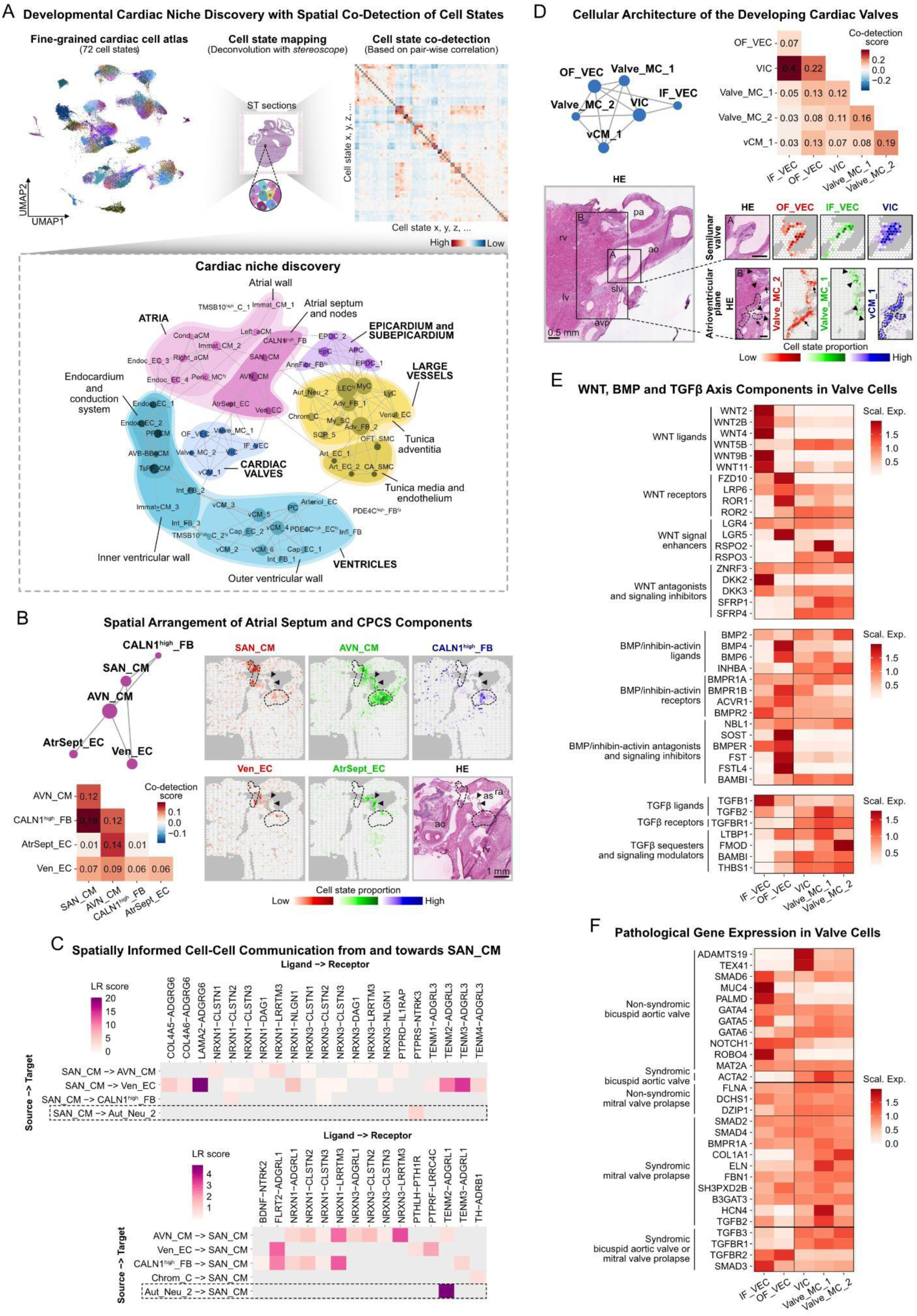
Spatially Informed Analysis of Developmental Cardiac Niches. **A**. Schematic representation of niche discovery strategy, using spatial mapping results of 72 fine-grained cell states for co-detection analysis (upper). Network graph of cardiac niches generated based on positive co-detection scores, with grey lines representing spatial association, and circle size reflecting the number of co-detected cell states (lower). **B.** Atrial septum and CPCS niche network graph and corresponding co-detection scores (left). Spatial mapping of SAN_CMs (red), AVN_CMs (green), CALN1^high^_FBs (blue) (upper right), Ven_ECs (red) and AtrSept_ECs (green) (lower right), outlining the atrial septum (arrowheads) and nodal tissue (dashed lines) in a 11 pcw heart section. Scale bar represents 1 mm. **C.** Spatially informed cell-cell communication analysis between SAN_CMs, its cellular neighborhood and Chrom_Cs, highlighting ligand-receptor interactions included in a recently published neural-GPCR module of CellPhoneDB^51^, differentially expressed in the analyzed cell states within their respective subsets. **D.** Cardiac valve niche network graph and corresponding co-detection scores (left). Spatial mapping of OF_VECs (red), IF_VECs (green), VICs (blue) (right, ROI A), and Valve_MC_2 (red), Valve_MC_1 (green) and vCM_1 (blue) (right, ROI B) cells, highlighting dominant cellular components of semilunar valves (ROI A) and neighboring atrioventricular plane regions (ROI B), respectively, in a 12 pcw section. Scale bars represent 0.5 mm. **E.** Heatmap representing per-gene scaled expression of WNT, BMP and TGFβ signaling molecules in valvular endothelial and mesenchymal cell states. **F.** Heatmap displaying per-gene scaled expression of bicuspid aortic valve disease– and mitral valve prolapse-related genes in valvular endothelial and mesenchymal cell states. In panel B-C: rv–right ventricle, lv–left ventricle, ra–right atrium, ao–aorta, pa–pulmonary artery, avp–atrioventricular plane, slv–semilunar valve, as–atrial septum, HE– hematoxylin-eosin.

Our analysis provided additional insight into the cellular composition and molecular interaction within the cardiac pacemaker-conduction system (CPCS). We found substantial overlap between predicted positions of CALN1^high^_FBs, SAN_CMs and AVN_CMs, providing further support to our assumption that CALN1^high^_FBs, in fact, represent a mesenchymal component of the developing nodal tissue. Of note, AVN_CMs showed the highest spatial co-detection scores with the AtrSept_EC state, implying the presence of conductive cardiomyocytes in the atrial septum of the fetal heart (Fig. 7B). We also identified several potential ligand-receptor pairs between SAN_CMs and their cellular surroundings (Figure 7C) using a recently published neural-GPCR module of CellPhoneDB^51^. These include multiple putative interactions directed from the pacemaker cells towards closely localized venous endothelial cells (Ven_EC), as well as molecular pairs likely involved in contact formation with autonomic neurons (Aut_Neu_2). Notably, CALN1^high^_FBs displayed a significant overlap in their interaction profile with SAN-CMs to a glial cell state recently discovered in the adult heart^51^, which suggests a potential hybrid function for this cell state in the developing heart. The ventricular CPCS components showed closest association with each other, and two ventricle-enriched endocardial cell states (Endoc_EC_1-2). We observed a temporal shift in the codetection of PF_CMs from Endoc_EC_1 towards Endo_EC_2, as well as differences in ligand-receptor interactions related to trabeculae formation and Purkinje fiber specification, highlighting functionally relevant molecular heterogeneity in the ventricular endocardium (Suppl. Fig. 7B).

Our analysis also demonstrated compositional differences between the great arteries (Art_EC_1, OFT_SMC, Adv_FB_1-2) and coronary arteries (Art_EC_2, CA_SMC, Adv_FB_2), consistent with differences in their developmental origins and functions, and a close spatial association between lymphatic endothelial (LEC^fg^) and myeloid cells (My_C), essential for *de novo* lymphangiogenesis (Suppl. Fig. 7C)^81^. In parallel, we observed a temporal decrease of co-detection scores between endothelial cell states in consecutive segments of the coronary vasculature (Art_EC_1-2, Arteriol_EC, Cap_EC_1-2), reflecting their gradual spatial separation (Suppl. Fig. 7D).

By identifying five, spatially distinct valvular endothelial and mesenchymal cell states, our study delineates the developing human cardiac valve architecture with unprecedented spatial and cellular resolution (Fig. 7D)^65,82^. Hence, we used this insight to map the distribution of WNT, BMP and TGFβ signaling network components, known to orchestrate valvulogenesis in a nuanced spatiotemporal manner, and also to be involved in the development of congenital and acquired valve disease (Fig. 7E)^83^. Notably, the endothelial populations showed the most distinct pattern of these transcripts, with IF_VECs displaying pronounced enrichment of WNT ligands (*WNT2*, *WNT2B*, *WNT4*, *WNT9B*, *WNT11*), antagonists and signaling inhibitors (*ZNRF3*, *DKK2*, *DKK3*), while OF_VECs showing higher expression of WNT receptors (*FZD10*, *ROR1*, *ROR2*) and signal enhancers (*LGR5*), besides certain BMP ligands (*BMP4, BMP6*) and antagonists (*SOST*, *FST*, *FSTL4*). Fluid forces, acting through the laminar flow-induced KLF2 (and KLF4) transcription factors, have been identified as major regulators of the activation of WNT-, and suppression of BMP-mediated signaling^72,84^. Accordingly, side-specific enrichment of WNT and BMP ligands in cardiac valves has been observed in several animal models, however, our study is the first to present consistent compartmentalization of various components of these signaling networks in developing human hearts. Regarding the TGFβ axis, cell state-related differences appeared less pronounced, with IF_VECs expressing higher levels of ligands (*TGFB1*, *TGFB2*), and OF_VECs of signaling modulators (*LTBP1*, *BAMBI*, *THBS1*). The valvular mesenchymal cell populations exhibited a more homogenous expression profile of these genes, including common enrichment of *WNT5B* and *RORA*, mediators of non-canonical WNT signaling, which has shown a positive correlation with the amount of calcification and fibrosis in diseased adult aortic valves^85^. Notably, Valve_MC_1 cells showed strong enrichment of *RSPO2*, implicated in the development of bicuspid aortic valve disease^86,87^, and Valve_MC_2 cells of *FMOD*, which has been proposed as a key molecule in the formation of a fibrous anchor area between cardiac valves and adjacent tissue segments^86^, consistent with the predominant localization of these clusters.

Cardiac valves are often affected in congenital and acquired heart diseases, however, the cellular culprits in these pathologies are often unclear. Thus, we explored the expression pattern of genes implicated in syndromic and non-syndromic forms of two major cardiac valve anomalies, bicuspid aortic valve disease (BAV) and mitral valve prolapse (MVP)^88–90^, across the valvular cell states (Fig. 7F). While most MVP-related genes, even in the non-syndromic form of the disease, appeared to have a broad distribution, many genes causing non-syndromic forms of BAV showed strong enrichment in IF_VECs and VICs, outlining these cell states as most susceptible to alterations of the analyzed genes. Our results showcase how future genetic studies on cardiac malformations could benefit from the spatially informed cell state decomposition of our dataset, providing support to potential gene-disease associations by presenting enrichment of pathological gene candidates in relevant developmental cell states.

## DISCUSSION

The majority of congenital, as well as several acquired heart diseases trace their origins to early development, highlighting the importance of this period in defining a healthy cardiac architecture. Exploring concurrent spatiotemporal cellular and molecular patterns can provide cues for the pathomechanisms, and thus potential therapeutic targets, in these conditions. Several human cardiac single-cell atlases were published in recent years, characterizing the cellular composition of developing, healthy and diseased adult hearts^23,91,92^. While representing invaluable resources about the cellular complexity of cardiogenesis, biological interpretation of the developmental datasets is often challenging due to their limited size or preselection of cell types of interest. Importantly, many of these approaches overlook the spatial context of the investigated cells, which holds key information during organ development. The possibility of combining single-cell analysis with unbiased, transcriptome-wide spatial gene expression information opened new horizons in understanding cardiac architecture^51^.

Our current work, built on the integrated analysis of single-cell and spatial transcriptomic datasets, presents the first comprehensive spatiotemporal atlas of heart development in the 1^st^ trimester. We demonstrate that, beyond independent exploration of regional expressional patterns, this approach also provides a basis for highly refined dissection of cell state heterogeneity, even in the heart’s minor structural or functional components. Accordingly, we performed detailed molecular characterization of specialized cardiomyocytes in the developing cardiac pacemaker-conduction system, highlighting their distinct electrophysiological properties and close spatial and functional association with specialized fibroblast cell states in the nodes, endocardial cells in the ventricular conduction system, and neurons in the developing autonomic innervation. Importantly, we present a resident neuroendocrine chromaffin cell population in the fetal human heart, with implications on organ-level cardiac response to hypoxia, and highlighting them as the potential cellular origin of the rare cases of cardiac pheochromocytoma. By resolving cellular complexity in cardiac valves, we identified several mesenchymal and endothelial populations in this location, including ectomesenchymal derivatives of the neural crest, and distinct endothelial cell states on opposite sides of the valves from as early on as the 6^th^ postconceptional week. We also explored the expression patterns of major signaling network components involved in valvulogenesis, reflecting flow-related transcriptomic differences, and highlighting endothelial cells on the ventricular side and valve interstitial cells as main cellular targets in non-syndromic bicuspid aortic valve disease. Furthermore, we identified a separate endocardial cushion-related population in the atrial septum with strong enrichment of the transcription factor *LEF1* and *MSX1*, lending support for a potential causative role of these gene in the development of atrial septal defects. The strong and temporally increasing enrichment of genes involved in endothelial-to-mesenchymal transition in this cell state is especially intriguing, considering the limited extension of the mesenchymal cap on the developing atrial septum, and the lack of persisting mesenchymal components in this structure, unlike in the cardiac valves. In lack of characteristic marker genes, we also leveraged spatial predictions on cell state distributions to untangle the diversity of non-mural cardiac fibroblast and mesenchymal cells in the developing heart, identifying a range of cell states associated with the annulus fibrosus, vessel adventitia, subepicardium, and myocardial interstitium. Moreover, we identified a fetal cardiac cell state resembling fibro-adipogenic progenitor (FAP) characteristics, which may constitute the developmental origin of cells responsible for fibro-fatty tissue transformation in arrhythmogenic right ventricular cardiomyopathy. Finally, we delineated the cellular composition of fine-grained cardiac niches in the developing heart, allowing for more nuanced and spatially informed interrogation of cell-cell communication and developmental trajectories, before bioinformatic tools, directly grounded in spatial transcriptomics data analysis, become established.

While this work captures the developing human cardiac architecture with unprecedented cellular and spatiotemporal resolution, it is intrinsically limited by the analyzed timeframe and size of the collected datasets. In fact, many genes implicated in the development of congenital heart diseases are active already during the first two weeks of cardiogenesis, which are not covered by our analysis. A larger sample size would allow for more robust interrogation of temporal gene expression changes even in less abundant cell states and could provide means to corroborate or even further refine cell state annotations presented in our work. Furthermore, balanced sampling of structural cardiac components for spatial transcriptomics analysis, especially in the case of morphologically not discernible structures like the nodal tissue, is challenging, which, however, can be mitigated by increasing the number of sections included in the analysis. Finally, the presented cellulo-architectural framework based on transcriptomics analysis could be enriched by a multiomics approach, mapping the proteomic, metabolomic, and chromatin accessibility landscape of the developing heart.

Further analysis of the presented dataset has the potential to furnish novel insight into other relevant questions of early cardiogenesis, not covered by our current study. The dataset can also be utilized as a spatiotemporal reference to assess expression patterns of CHD candidate genes in the 1^st^ trimester, and for iPSC-derived cardiac cell and tissue therapies, often resembling embryonic phenotypes. In conclusion, our work underscores the vast and yet underexploited possibilities of spatially targeted molecular analysis within cardiac research.

## METHODS

### Collection of Human Developmental Heart Tissue Samples

All heart specimens included in this study were collected from elective medical abortions at the Department of Obstetrics and Gynecology at Danderyd Hospital and Karolinska Huddinge Hospital in Stockholm, Sweden. All patients were over 18 years old and donated tissue with written informed consent, after receiving both oral and written information about the research project, and the possibility of retracting their consent at any time, including later destruction of the donated tissue. Following collection, the specimens were transferred from the clinic to the dissection laboratory. Heart tissue samples were then swiftly dissected under sterile conditions within 2 hours post-abortion. In total, 36 human developmental hearts between the age of 5.5 and 14 postconceptional weeks (pcw) were included in this study. The embryonal/fetal age was determined using clinical data (ultrasound and last menstrual period), anatomical landmarks and actual crown-rump-length. Both sexes were present in the full dataset, specifically 17 hearts were from female and 14 from male donors, while information on sex for the 5 hearts used for *in situ* sequencing and immunofluorescence experiments was not obtained. The study was performed with approval of the Swedish Ethical Review Authority and the National Board of Health and Welfare, under the ethical permit number 2018/769-31.

### Sample Preparation for Spatial Methods

In total, 21 collected heart samples were embedded in Tissue-Tek O.C.T., snap-frozen, and stored at –80 °C. Samples were cryosectioned at 10 µm thickness. Sections from 16 hearts were placed on Visium Gene Expression glass slides, while sections from 5 other hearts were placed on Superfrost glass slides for *in situ* sequencing and immunohistochemistry.

### Visium Spatial Gene Expression Library Preparation and Sequencing

Spatial gene expression libraries were generated using the 10x Genomics Visium Gene Expression kit, according to the manufacturer’s protocol^93^. A minimum of two consecutive tissue sections were included in the analysis as technical replicates for all but one heart sample, where replicates had to be excluded from downstream analysis due to experimental deficiencies. Tissue images were taken at 20x magnification using the Metafer Slide Scanning platform (Microscope: AxioImager.Z2 with ScopeLED Illumination, Zeiss; Camera: CoolCube 4m, MetaSystems; Objective: Plan-Apochromat 20X/0.80 M27, Zeiss; Software: Metafer5). The permeabilization time was adjusted to 20 minutes. Raw images were stitched with VSlide software (MetaSystems). In total, 38 Visium libraries were prepared from 16 hearts. Libraries were sequenced by using Nextseq2000 (Illumina), where length of read 1 was 28 bp and read 2 was 120 bp long.

### Processing and Analysis of Visium Spatial Gene Expression Data

Sequenced libraries were processed using Space Ranger (v.1.2.1; 10x Genomics). Reads were aligned to built-in human reference genome (GRCh38 v.2020-A, Ensembl 98). Further processing and data analysis of the spatial data was performed in R Statistical Software (v.4.0.5)^94^ using STUtility (v.0.1.0)^95^ and Seurat (v.4.1.1)^96^ packages. The created count matrix was filtered for MALAT1, ribosomal, mitochondrial, and hemoglobin genes. Spots with fewer than 200 genes were removed from downstream analysis. The processed data was described with a median of nFeature_Spatial and nCount_Spatial of 2,191.5 and 4,179, respectively. Each tissue section was normalized separately using SCTransform function from the Seurat package. Sections with at least 3 heart chambers present were selected for clustering. In the following steps, principal component analysis (PCA) and sample integration using Harmony (v.1.0)^97^ was performed. Unsupervised clustering of spots was performed using the shared nearest neighbor (SNN) algorithm from the Seurat package. Uniform manifold approximation and projection (UMAP) was used for cluster visualization. Temporal evolution of spatial clusters was analyzed by embedding the original UMAP with three age groups (age 1: 6-7 pcw; age 2: 8-9 pcw; age 3: 10-12 pcw). Downsampling in age groups 2 and 3 was performed to equalize the spot numbers across embeddings (n = 2,649). FindAllMarkers() function from Seurat package (logfc.threshold = 0.5, only.pos = TRUE, min.pct. = 0.01) was applied to identify differentially expressed genes (DEGs) between clusters. Subsequently, all clusters were manually annotated. Spatial feature plots were generated with the STUtility package.

In addition to standard clustering, region segmentation was performed on the same subset of sections using the R tool Banksy (v.1.0.0)^41^. This spatially-aware clustering enables deciphering neighborhoods using both the transcription information of a spot and its neighbors. The hyperparameter k was set to 6 to select the first layer of direct surrounding Visium spots. The second hyperparameter, lambda, was set to 0.8 to focus the algorithm to focus on region segmentation, as it weighs the importance of the spatial component for the clustering. Banksy neighborhood-augmented feature space’s clustering was performed with the default Leiden algorithm at a resolution of 0.9. Resulting regions were transferred to the existing object for exploration and visualization.

Non-negative matrix factorisation (NMF) was performed jointly using the singlet package (v.0.99.36)^98^ in R implemented in the semla toolkit on the entire Visium dataset. The parameter k was manually fixed to 20, resulting in 20 shared factors across the dataset. This widely used method in the single-cell and spatial omics field decomposes data variability into gene modules that covariate in a latent space. Each factor consists of all the initial variable features ranked by order of influence in the factor-forming feature loadings.

Hematoxylin-eosin micrographs of the analyzed tissue sections were included in the figures after linear adjustments of intensity and contrast, performed in Affinity Designer. In composite figures with micrographs of more than one tissue section, orientation of the images was adjusted according to the anatomical position (left-right, superior-inferior) of the sampled cardiac structural components.

The original micrographs, quality metrics, sex and age distribution of the heart sections included in the Visium dataset are displayed in Extended Figure 8A-D. Batch effects in the data were visually assessed in the section selection used for clustering analysis, by embedding the UMAP with the individual samples, either in an integrated or in an age-resolved manner (3 age groups) (Ext. Fig. 10A-B). Furthermore, cluster and region distributions were computed per section and represented as barcharts enabling an evaluation of consistency between technical replicates (Ext. Fig. 11A-B). Similarly, coarse-grained cell type proportions were computed per section for the entire dataset and visualized in barcharts (Ext. Fig. 11C).

### *In Situ* Sequencing

To generate the *in situ* sequencing (ISS) datasets presented in the manuscript, a panel of 150 gene targets was created, including known regulators of cardiogenesis, consensus markers of major cardiac cell types, and highly enriched genes of clusters of interest (Ext. Tab. 1). For these genes, a total of 5 padlock probes per gene were designed, following a pipeline described by Lee et al.^99^. Next, a direct RNA *in situ* sequencing protocol^99^ was applied to 9 tissue sections, corresponding to 6.5 (section_1-3), 8.5 (section_4), 9 (section_5-6) and 11.5 pcw (section_7-9) (Ext. Fig. 1e).

For ISS sequencing library preparation, the sections were thawed at RT for 5 minutes, washed with PBS, and fixed in 3% paraformaldehyde for 5 minutes. The sections were then washed 3x in PBS, permeabilized with a 0.1M HCl solution for 5 minutes, and washed 3x in PBS. Next, the sections were sequentially dehydrated in a 70% and 100% ethanol bath for 2 minutes each. Secure-seal chambers (-SA50, 1-13mm Diameter X 0.8mm Depth, 22mm X 25mm OD, 1.5mm Diameter Ports) were laid over each sample, which were then rehydrated with 0.5% Tween-PBS, followed by a PBS wash. A padlock probe solution (2x SSC, 10% formamide, 10 nM of each padlock probe) was added to the chambers for overnight incubation in a humidified chamber at 37 °C. The sections were then washed 2x with a washing solution (2x SSC, 10% formamide), and then 2x with 2x SSC. The sections were then incubated in a ligation mix (1x T4 Rnl2 reaction buffer (NEB, B0239SVIAL), 0.05 µM RCA primer, 1 U/µl RiboProtect (BLIRT, RT35), 1.0 U/µl T4 Rnl2 (NEB, M0239)) in a humidified chamber at 37 °C for 2 hours. The sections where then washed 2x with PBS, and then incubated in a rolling circle amplification mix (1x phi29 buffer (50mM Tris-HCL pH 8.3, 10mM MgCl2, 10mM (NH4)SO4), 5% glycerol, 0.25 mM dNTPs (BLIRT, RP65), 0.2 µg/ml BSA and 1U/µl Φ29 polymerase (Monserate Biotech, 4002)) at 30 °C in a humidified chamber overnight.

After washing the sections 3x with PBS and 2x with 2x SSC, fluorescent probes were hybridized to the samples with incubation in a bridge probe mix (20% formamide, 2x SSC, 0.1 µM of each bridge probe) for 30 minutes in a humidified chamber at RT. The sections were washed 2x with PBS and 2x with 2x SSC, followed by hybridizing a detection oligo mix (20% formamide, 2x SSC, 0.5µM of each detection oligo (conjugated with Atto425, AlexaFluor488, Cy3, Cy5, AlexaFluor750), 0.25 µM DAPI (Biotium, S36936)) for 30 minutes in a humidified chamber at RT. Sections were washed 2x with PBS, then mounted with SlowFade Gold Antifade Mountant (Thermo Scientific, S36936).

Cyclical imaging of the sections^99^ was performed by stripping bridge probes and detection oligos after each cycle with 3x washes in 100% formamide for 5 minutes each, followed by 5x washes in 2x SSC. A new set of bridge probes were then hybridized in the same manner, and the process was repeated until all 5 cycles were imaged. Image acquisition was performed using a Leica epifluorescence microscope (Microscope: DMi6000, Lumencor® SPECTRA X Light Engine, LMT200-HS automatic multi-slide stage; Camera: sCMOS, 2048×2048 resolution, 16 bit; Objective: HC Plan-Apochromat 20X/0.80 air, and Plan-Apochromat 40X/1.10 W CORR oil objectives; Filters: filter cubes 38HE, Chroma 89402 – ET – 391-32/479-33/554-24/638-31 Multi LED set, Chroma 89403 – ET – 436-28/506-21/578-24/730-40 Multi LED set, and an external filter wheel DFT51011). Each region of interest (ROI) was marked and saved in the Leica LASX software for repeated imaging. Each ROI was automatically subdivided into tiles, and a z-stack with an interval of 0.5 µm was acquired for each tile in all the channels, with a 10% overlap between each tile. The ROIs were saved as TIFF files with associated metadata.

To create the *in situ* sequencing maps^99^ featured in this manuscript, the images obtained for each dataset across cycles were orthogonally projected using maximum intensity projection with the aim of collapsing the different z-planes into a single image per tile. Next, tiles from individual cycles were stitched, and images from different cycles were aligned using ashlar, obtaining 5 large, aligned TIFF files for each dataset, corresponding to the different imaging rounds. Due to computational considerations, images were then retiled into smaller aligned images. A pre-trained CARE model was then applied to each of the retiled images, resulting in a reduction of the point spread function and increasing signal to noise ratio. The output was then converted to the SpaceTx format and spots were identified using the PerRoundMaxChannel decoder, included in starfish. Every spot was given a quality score for each of the cycles, by normalizing for the most intense channel. Using the quality scores from all rounds, two quality metrics were composed for each spot: the mean quality score, defined as the mean of the qualities across cycles, and the minimum quality score, which corresponds to the score of the round where the quality was the lowest. Using these metrics, spots were filtered, retaining only the ones presenting a minimum quality score over 0.4 for further analysis. Finally, using the identity and position of all the high-quality score spots decoded, expression maps were assembled for each of the datasets.

For ISS panels included in the figures, visualization of selected target genes over the DAPI image of the tissue section was performed using TissUUmaps^100^, with high resolution capture of the image viewport (zoom factor=8). Orientation of the images was adjusted according to the anatomical position (left-right, superior-inferior) of the sampled cardiac structural components.

Consistency between technical replicates in the ISS dataset was evaluated based on agreement between detected transcript numbers of individual targets, by calculating Pearson correlation coefficient between paired sections (Ext. Fig. 11D).

### Indirect Immunofluorescence and Imaging

After fixation with 4% paraformaldehyde (Thermo Scientific, 043368.9M) for 15 minutes, the heart sections were washed 3x in PBS for 15 minutes. Then, the samples were incubated overnight at 4°C with the following antibodies: anti-ARL13B (RRID:AB_3073658, ab136648, Abcam, 1:400), anti-PCNT (RRID:AB_2160664, ab28144, Abcam, 1:800) and anti-PDE4C (RRID:AB_3094595, HPA054218, Atlas Antibodies, 1:100). The sections were then blocked for 30 minutes in 1xTBS (Medicago, 097500100) with 0.5% TSA blocking reagent (Perkin Elmer, FP1020) and Hoechst (Invitrogen, H3570, 2μg/mL). Next, the samples were incubated for 90 minutes at RT with anti-mouse-IgG2a-Alexa647 (RRID:AB_2535810, A21241, Invitrogen, 1:800), anti-mouse-IgG1-Alexa555 (RRID:AB_2535769, A21127, Invitrogen, 1:800) and anti-rabbit-A488 (RRID:AB_2576217, A11034, Invitrogen, 1:800) secondary antibodies, diluted in the blocking buffer. After incubation, the slides were washed 3x with TBS-Tween for 15 minutes. Coverslips (VWR, 631-0147) were mounted on the slides using Fluoroshield mounting medium (Invitrogen, 00495802). Imaging was performed with a PhenoImager Fusion 2.0 scanner, using a 20x objective and a phenocycler fresh frozen filter set. Exposure times were 5 ms for Hoechst (DAPI channel), 300 ms for Alexa488 (FITC channel), 250 ms for Alexa555 (ATTO550 channel), and 400 ms for Alexa647 (Cy5 channel). PhenoImager automated algorithm was used for the selection of focal points.

The captured tiles were automatically stitched together by the software operating PhenoImager, which outputs a whole-slide image in the QPTIFF format. The QPTIFF files were opened in QuPath^101^. A rectangular ROI was selected that circumscribed the whole imaged tissue. The content of the ROI was sent to an ImageJ distribution in QuPath (without changing scale or intensity values) and the image was saved as a TIFF file. Next the TIFF file was opened in FIJI. Linear adjustments of intensity and contrast were performed separately for each detection channel on the collected images. Individual zoom-in ROIs were selected and the regions in the ROIs were duplicated and saved as png files for implementing them into the figure. To show the whole section in the figure, the whole image was downscaled in FIJI to 25% of the original size and exported as a png file for embedding into the figure.

### Sample Preparation for Single-Cell RNA Sequencing

In total, 15 heart samples were analyzed by single-cell RNA sequencing. The specimens were cut into smaller pieces and minced using a blade. The tissue was transferred into a 15 ml Falcon tube containing an enzymatic solution of Collagenase II (200 U/ml, Worthington Biochemical, LS004174), DNase I (1 mg/ml, Worthington Biochemical, LK003170) in Earle’s balanced salt solution (EBSS, Worthington Biochemical, LK003188) which had been pre-oxygenated with 95% O2:5% CO2 on ice for 5-10 minutes. Tubes were put in a water bath and the samples were digested at 37 °C with an incubation time ranging from approximately 45 minutes to over 2.5 hours, depending on the age of the tissue (tissues from later developmental stages required longer dissociation times). During the incubation, the cell suspension was triturated and dissociated around every 25 minutes, using glass fire-polished Pasteur pipettes, with gradually decreasing tip diameter for each round to enhance dissociation. The cell suspension was manually inspected under the microscope to assess the level of digestion, followed by a filtering step using a prewet 30 μm cell strainer (CellTrics, Sysmex) to remove debris, fibers and smaller undissociated remains. A small volume of EBSS was added to the filtered suspension to dilute and inactivate the enzyme, then cells were pelleted through centrifugation at 200 g for 5 minutes. If needed, gradient centrifugation was performed to remove blood cells. For this step, the cell pellet was resuspended in a solution containing 900 μl EBSS, 100 μl albumin inhibitor solution (Ovomucoid protease inhibitor and bovine serum albumin, OI-BSA, LK003182) reconstituted in EBSS, and 50 μl of DNase I (larger volumes with the same reagent concentrations were used in case of higher cell numbers). The resuspended solution was carefully layered on top of 3 ml albumin inhibitor solution in a 15 ml Falcon tube and gradient centrifugation was performed at 70 g for 4 to 6 minutes. Then the supernatant was removed, and the cells were resuspended in a minimum amount of EBSS, starting from 100 μl (or more in case of higher cell numbers). The cell suspension was transferred to a 1.5 ml Eppendorf tube, pre-coated with 30 % BSA (A9576, Sigma-Aldrich). If a few cellular clumps had formed, a small volume of BSA was added to the suspension (starting with 0.3 % BSA of the total volume). Cells were counted on a hemocytometer (Bürker/Neubauer chamber) and cell viability was assessed with Trypan blue (Gibco, 15250061). Cell concentrations were adjusted to values optimal for the 10x Genomics Chromium analysis.

### Single-Cell RNA Sequencing of Human Embryonic and Fetal Heart Cells

Single cells were captured using the droplet-based platform Chromium (10x Genomics), and the Chromium Single Cell 3′ Reagent Kit v2 and v3. Due to a change in the chemistry suppliance of the kits, 5 out of 21 libraries were sampled with v2. Cell concentrations were adjusted to between 800-1200 cells/μl where possible, for a target recovery of 5000 cells per library. Cells were loaded on the Chromium controller and downstream procedures such as cDNA synthesis, library preparation and sequencing were done according to the manufacturer’s instructions (10x Genomics). Libraries were sequenced between approximately 100,000 to 250,000 reads/cell on the Illumina NovaSeq platform, reaching a saturation between 60 to 90%. All libraries were demultiplexed using Cell Ranger (cellranger mkfastq v.4.0.0; 10x Genomics) and filtered through the index-hopping-filter tool (v.1.1.0) by 10x Genomics. STARSolo (v.2.7.10a) was used to determine Unique Molecular Identifier (UMI) counts, with the following parameters used:

--soloFeatures Gene Velocyto

--soloBarcodeReadLength 0

--soloType CB_UMI_Simple

--soloCellFilter EmptyDrops_CR %s 0.99 10 45000 90000 500 0.01 20000 0.01 10000

--soloCBmatchWLtype 1MM_multi_Nbase_pseudocounts

--soloUMIfiltering MultiGeneUMI_CR

--soloUMIdedup 1MM_CR

--clipAdapterType CellRanger4

--outFilterScoreMin 30

Reads were aligned using the GRCh38.p13 gencode V35 primary sequence assembly. To minimize the loss of reads that occur when they map to more than one gene, we applied a read-filtering step with certain criteria, which is described in detail, along with the retrieval of count matrices and read alignments, in Braun et al.^102^.

### Processing and Filtering of the Single-Cell RNA Sequencing Data

Further processing and analysis of the scRNA-seq data was performed in R (v.4.3.1)^94^, using the Seurat toolkit (v.4.3.0.1)^96^. The data was jointly handled as one Seurat object, and an initial filtering step was applied, removing cells with more than 30% mitochondrial transcript counts. Cells with less than 250 unique genes or less than 3% ribosomal protein-coding transcript content were also removed unless they also had more than 10% hemoglobin gene expression. The latter step was included to keep red blood cells for the subsequent doublet detection step. From the 107,673 initial cells imported, 101,104 cells remained after the first filtering round.

Doublet detection was performed separately for each sequencing dataset using the DoubletFinder R package (v.2.0.3)^103^, with expected multiple rates defined according to 10x Genomics guidelines. The data was then merged again, and a second filtering step was applied, removing expected doublets (4,568 doublets), as well as cells with more than 10% hemoglobin gene expression (19,742 red blood cells), identifying 76,991 high-quality cells. Data was then scaled using the Seurat function ScaleData() based on the top 4,000 most highly variable genes, regressing out the number of reads, number of unique genes, percentage of ribosomal genes, percentage of mitochondrial genes, percentage of hemoglobin genes, percentage of heat shock protein-related genes, as well as S and G2M scores. Dataset integration was performed using the R package Harmony (v.0.1.1)^97^, leveraging the top 50 principal components. The processed data was described with a median of nFeature_RNA and nCount_RNA of 2,417 and 4,838, respectively.

Quality metrics, sex and age distribution of the heart samples included in the single-cell RNA sequencing dataset are displayed in Extended Figure 12A-B.

### Age Categorization for Single Cell Data Analysis

To facilitate a temporal analysis of the single-cell RNA sequencing dataset, samples were classified into four age groups: age 1 (5.5-6 pcw), age 2 (7-8 pcw), age 3 (9-11 pcw), and age 4 (12-14 pcw). This categorization enabled a comprehensive examination of temporal trends within the data, providing insights into developmental patterns across different developmental stages.

### Coarse-Grained Clustering and Analysis of the Single-Cell RNA Sequencing Data

Coarse-grained clusters (referred to as HL, as in ‘high level’, in the code) were obtained using the Louvain community detection algorithm on an SNN graph built from the harmony embedding (nearest neighbors set to 20) with the resolution parameter set to 1.8. The UMAP embedding was computed from the harmony embedding (top 50 components), with 10 neighbors and 100 epochs. To identify differentially expressed genes and facilitate the computation, the clusters were downsampled to have a maximum of 150 cells, and pair-wise comparison of each cluster against all other clusters was performed using the FindAllMarkers() function from Seurat (max.cells.per.ident = 300, logfc.threshold = 0.1, min.pct. = 0.05). To maintain the highest heterogeneity within each cluster, the x=150 cells per group were randomly picked with the R function sample(colnames(data)[data$clusters_louvain_oi == x], size = sample_size[x]), without using any QC condition. Further subclustering of the mixed coarse-grained cluster 18 was performed with the same settings and principal components, except for using the 10 nearest neighbors and a resolution of 0.25, resulting in two distinct subclusters. The small coarse-grained cluster 35 was merged with the highly similar cluster 4 for further analysis.

For temporal analysis of the coarse-grained cluster distribution, UMAP embedding was performed across the four predetermined age groups. Downsampling in age groups 1, 2 and 4 was performed to equalize the cell numbers across embeddings (n = 8,742).

The original coarse-grained clusters and related quality metrics are displayed in Ext. Fig. 12C-E. Coarse grained cluster 6 and 34 marking red blood cells, and 19 and 27 featuring low quality cells were excluded from downstream analysis (Ext. Fig. 12F). Batch effects in our data were visually assessed by embedding the coarse-grained UMAP with individual samples, either in an integrated or in an age-resolved manner (4 age groups) (Ext. Fig. 10A, C).

### Fine-Grained Clustering and Analysis of the Single-Cell RNA Sequencing Data

To create fine-grained clusters (occasionally referred to as DL, as in ‘deep level’, in the code), cardiomyocyte (referred to as CM in the code), endothelial (referred to as EN in the code), mesenchymal cell-fibroblast (referred to as FB in the code), and innervation-related cell (referred to as IN in the code) subsets were all processed similarly. The subsetted data was again scaled, this time based on the top 3,000 highly variable genes, and the same variables were regressed out as for the complete dataset. Again, the data was integrated using Harmony, based on the top 50 principal components for the specific subset. Clustering was based on the 30 nearest neighbors and calculated using Louvain community detection with a resolution of 1.35 for cardiomyocytes, 1.25 for endothelial cells, 1.5 for fibroblasts, and 1 for innervation-related cells. UMAPs were calculated based on the top 50 integrated principal components and 30 neighbors, using 100 epochs. Marker genes were identified using the FindAllMarkers() from Seurat, using the same settings as for the coarse-grained clusters. The pacemaker-conduction system-related cardiomyocyte clusters 4, 8, and 19 were further subclustered to increase granularity. For this purpose, Louvain community detection was applied to each of these clusters, calculated from the top 30 nearest neighbors with a resolution of 0.5.

Fine-grained cardiomyocyte clusters 11, 13, 15, 16, 20, 21, and 22, fine-grained endothelial clusters 4, 8, 9, and 22, fine-grained mesenchymal cell-fibroblast clusters 10, 16, 21 and 22, and fine-grained innervation-related cluster 2, 5, 6, 9, 10, 14, 15, 16 and 18 were excluded from downstream analysis, on the basis of low quality metrics, potential doublet contamination, or lack of consensus marker expression of the relevant cell types. The original UMAPs of these cell subsets, consensus marker expression profile of original fine-grained clusters, and parameters used as basis for cluster exclusion are presented in Extended Figure 13A-E.

### Spatial Mapping Using Cell State Deconvolution

The spatial mapping of single-cell transcriptomics data onto tissue sections included in the Visium dataset was achieved through probabilistic inference using *stereoscope* (v.0.3.1)^104^, effectively addressing spatial heterogeneity. By assuming a negative binomial distribution, this model estimates the proportions of single-cell states at every spatial spot. This guided decomposition process was executed separately for coarse– and fine-grained annotations. In the coarse-grained iteration, with 31 clusters remaining after excluding poor-quality clusters and red blood cells, the dataset was downsampled to the 500 cells containing the highest number of features in each population. Then, the top 5,000 most variable genes from this subset were utilized. Similarly, low-quality clusters were excluded in the fine-grained iteration and the dataset was downsampled to the top 250 cells with the highest number of features for each of the 72 remaining cell states. The top 5,000 most variable genes from this subset were employed for the analysis. The *stereoscope* analysis ran with a batch size of 2,048 and 50,000 epochs for parameter estimation and proportion inference steps in both the coarse– and fine-grained iteration. A complementary, strictly age-matched cell type mapping was performed using ST and coarse-grained sc-RNA-seq data of the same age window. After excluding single-cell data from hearts not matching the age range of the ST sections, deconvolution with *stereoscope* was executed for the 3 previously determined age groups (age 1: 6-7.5 pcw; age 2: 8-9 pcw; age 3: 10-12 pcw) with 500 cells per population, the top 5,000 most variable genes, and unchanged *stereoscope* parameters.

Spatial cell state prediction maps were generated using MapFeatures() from the semla package (v.1.1.6)^105^. A max_cutoff parameter of 0.99 was applied to mitigate potential biases from outliers in coarse-grained clusters, while no max_cutoff was used for fine-grained clusters. In the figures, these maps were visualized alongside the corresponding intensity– and contrast-adjusted hematoxylin-eosin micrograph of the relevant tissue section.

### Annotation of Spatial and Single-Cell Clusters

Annotation of the spatial clusters was performed manually, based on the relative enrichment of consensus markers of cell states with characteristic spatial distributions, and the relation between known anatomical landmarks and the observed localization of spatial clusters within the analyzed heart sections. For the coarse– and fine-grained single-cell clusters, a spatially-aware annotation strategy was utilized. Cell state identification was based on the hierarchy of three features: first, the relative enrichment of consensus cell type/state markers was considered, discerning major cell populations and minor cell states with known transcriptomic profiles. For further refinement, the deconvolution results of the integrated single-cell and spatial transcriptomics datasets were utilized, factoring in the predicted localization of the analyzed cell states. Finally, enrichment of gene signatures consistent with different biological states in relation to maturation, metabolism, or proliferation were considered, in case the previous two features alone did not sufficiently explain the distinct characteristics of the cluster. Where possible, cluster names were selected to match the conventional cell state nomenclature, while in other cases, they were formulated to reflect the distinguishing spatial or transcriptomic feature of the population in question. Fine-grained cell states with annotations identical to coarse-grained clusters are distinguished by fg (as in ‘fine-grained’) superscript.

Dominant marker genes of single cell clusters were identified by sequential filtering of the top 40 DEGs by lowest p-values (p_val), top 20 DEGs with the highest percentage difference (pct.1-pct.2), and top 5 or 10 DEGs with the highest average log2 fold change (avg. log2FC), which were then visualized by dot plots across the compared clusters.

### Assessment of Ion Channel Profiles of Cardiac Pacemaker-Conduction System-Related Cell States

After differential gene expression analysis across all fine-grained cardiomyocyte clusters, a comprehensive list of ion channel-encoding DEGs (p < 0.05, avg. log2FC > 0), enriched in any of the fine-grained cardiomyocyte clusters, was extracted. Next, this list was filtered in parallel for genes expressed in more than 10% of cells (pct. exp. > 0.1) of the SAN_CM or AVN_CM clusters, or of the AVB-BB_CM, PF_CM or TsPF_CM clusters, respectively. In Fig. 3F, genes with higher average expression (avg. exp.) observed in SAN_CMs or AVN_CMs compared to the contractile atrial clusters Right_aCM, Left_aCM and Cond_aCM were included. In Suppl. Fig. 3F, genes with higher average expression (avg. exp.) observed in AVB-BB_CMs, PF_CMs or TsPF_CMs compared to the contractile ventricular cluster vCM_4 were included. The compiled gene lists were then assessed for biological relevance in the specification of the pacemaker-conductive phenotype.

### Analysis of Cell State Co-Detection and Identification of Spatial Compartments and Cellular Niches

Spatial compartments and cellular niches were identified based on co-detection scores, obtained by calculating the commonly used Pearson correlation coefficients between the distribution of each fine-grained cell state pair, based on the spatial cell proportion predictions of the stereoscope analysis. The matrix of these in-pair calculated correlation scores, referred to as co-detection scores, served as a proxy for appreciating the level of spatial overlap between the fine-grained cell states despite the sparsity of the data, outlining groups of cell states with close spatial association. Co-detection scores in our dataset ranged between –0.25 and 0.55 (95th percentile = 0.10) in the age-resolved, and between –0.2 and 0.42 (95th percentile = 0.09) in the merged dataset. The co-detection matrices were further visualized on heatmaps using geom_tile() from ggplot2 (v.3.4.4)^106^. Spatially co-localised cell types were described with positive co-detection scores, while non-co-localised cell types were described with negative ones. Since 72 cell types were used for the deconvolution, inherently many proportions were close to zero, hence, the co-detection heatmap resulted in many scores around zero. Those indicate neither co-detection, nor non-co-detection. To capture niches clearly, a co-detection graph network was built, where co-detection scores above a manually selected arbitrary threshold of 0.07 were kept. All the other scores were set to 0 since we considered building graphs with negative values not meaningful. From the remaining co-detection matrix, we used graph_from_adjacency_matrix() function from the igraph (v.1.5.1)^107^ package in R (weighted=TRUE, mode=“undirected”, diag=FALSE) which represents high co-detection scores by shorter edges between nodes (nodes appear closer in the graph) and degree() function from the igraph package in R to calculate the degree for each node influencing the size of the nodes (the more connections, the bigger the node). The graph was plotted using the plot() function from igraph package (vertex.size=deg*1.1) and other parameters left to default. In our case (less than 1000 vertices, and no other attributes), the network layout has been selected by default by the tool, resulting in the Fruchterman-Reingold algorithm. The resulting network (Figure 7A, lower panel) is useful for readability, where niches and compartments were manually annotated based on the cell states’ spatial relation to known cardiac structural components. These steps were performed on the merged Visium dataset, as well as in an age-resolved manner (age 1: 6-7.5 pcw; age 2: 8-9 pcw; age 3: 10-12 pcw) to describe cellular spatial relations in a temporal context.

Spatial association of manually selected niches was presented with a new in-house function CellCol.R, an adaptation of the MapFeatures() function built upon the package semla (v.1.1.6)^105^, enabling simultaneous visualization of per-spot predicted cell type proportion of two or three cell states, overlaid onto the section of interest in the Visium dataset. Overlap of co-detected cell states was presented through a scaled color blending using blend_colors() function from colorjam (v.0.0.26.900)^108^.

### Cell-Cell Communication Analysis

Single-cell clusters in spatial proximity were selected for cell-cell communication (CCC) analysis. Intercellular communication between ligands (L) and receptors (R) was inferred using CellPhoneDB (v.2)^109^ implementation through the tool provider Liana (v.0.1.13)^110^ in R (v.4.3.1). A mean of the gene expression for known ligand and receptor pairs (LR) from the curated CellPhoneDB was calculated by randomly permuting (100 epochs) cell labels in the subset, generating a null distribution for each LR event between each cluster in the subset. The p-value of 0.05 was used as threshold for selecting significantly enriched LR between cell states having a mean expression of LR equal or higher than the mean of that specific LR event calculated with the null distribution. The default mode *magnitude* within the function rank_method() was used to order LR events based on expression level. A selected subset of the identified ligand-receptor interactions was finally visualized as a dot plot across the analyzed cell state pairs.

Additionally, a subset of co-localised cell types selected from the niche network (SAN_CM and its direct neighbors and Chrom_C) accounted for the presence of the LR interaction described in a recently published, custom neural-GPCR module of CellPhoneDB (41586_2023_6311_MOESM5_ESM)^51^. After selecting the pairs of protein_name_a (ligands) and protein_name_b (receptors) present in the module, separate lists of ligands and receptors were generated for every cluster of interest from their differentially expressed genes. The cell-cell interactions were evaluated bi-directionally, SAN_CM (donor) to its neighbors (receivers) and from neighbors (donors) to SAN_CM (receiver), using the gene lists accordingly. The interactions were scored for each present LR pair by multiplying the average expression of the ligand and the receptor from their respective cell types with the AverageExpression(group.by = c(“annot_dl”)) function from the Seurat package. The results are represented as a heatmap where LR pairs present are colored according to their score, while a gray square is presented for LR pairs where the ligand or receptor was not enriched in the assessed cell states within their respective cellular subsets.

Spatial LR interactions were visually presented in tissue sections for a selection of known LR pairs by calculating their geometric means per spot. The spatial LR interaction scores were plotted using an in-house adaptation of the MapFeatures() function from the semla package.

### RNA Velocity and Pseudotime Analysis

RNA velocity for innervation-related fine-grained clusters was performed on the top 5,000 variable features calculated with filter_genes_dispersion() from scVelo (v.0.3.0)^111^ in Python (v.3.8.17). PCA was performed with Scanpy (v.1.9.5)^112^, followed by recomputing the neighboring graph (scanpy.pp.neighbors(), n_neighbors = 25). Velocity was plotted on the PCA embeddings after running scv.tl.velocity() and scv.tl.velocity_graph(). Pseudotime was calculated separately on the top 5,000 variable features (highly_variable_genes()) with scFates (v.1.0.6)^113^ in Python (v.3.11.6). PCA and clustering were performed similarly as for the RNA velocity analysis, resulting in the same embedding. Curves of the PCA were calculated using the ElPiGraph algorithm implemented in scFates (scFates.tl.curve()). Subsequently, pseudotime was calculated with scFates.tl.pseudotime() (n_jobs = 20, n_map=100), and both pseudotime and expression of markers were visualized on the PCA embedding (scanpy.pl.pca()).

### Transcription Factor Inference Using Gene Regulatory Networks

Inference of transcription factor (TF) activity was performed using the Scenic pipeline (v.1.3.1)^114^ in R (v.4.3.1). Gene regulatory networks (GRN) were generated based on co-expression across the single-cell RNA sequencing data with GENIE3 (v.1.22.0)^115^, based on the overlap between the most variable features calculated during clustering of the dataset and the provided databases (hg19-500bp-upstream-10species.mc9nr.feather and hg19-tss-centered-10kb-7species.mc9nr.feather). Co-expression of the remaining TFs in the network were converted into regulons using runSCENIC_1_coexNetwork2modules(). The implemented AUCell from Scenic was used to score every regulon in each individual cell. Then, known TF-binding motifs were used to prune interactions from the GRN (regulatory modules or regulons). Relevant regulons were retrieved if their specific TF was differentially expressed (FindMarkers(), test.use = “wilcox”, avg_log2FC > 0 and p_val < 5e-7) in the cluster of interest, compared to the other clusters of the specific group. Regulated genes in the dataset were similarly selected (FindMarkers(), test.use = “wilcox”, top 1,000 based on p-value). The resulting network of regulatory TFs and their associated target genes was constructed as a network graph using the igraph package^107^.

The entire list of inferred transcription factors and target gene pairs, as well as differentially expressed transcription factors in the investigated cell states are presented in Extended Table 2-9.

Regulon enrichment analysis in selected endothelial cell states was performed for each cell using AddModuleScore() from Seurat, scoring the average expression of the target genes of a TF compared to control sets of genes. The enrichments were visualized on violin plots for the three selected clusters in an age-resolved manner, where age groups 3 and 4 were merged (creating an age group representing 9-14 pcw) in order to decrease the difference in cell numbers between the compared populations.

### Pathological gene sets enrichment analysis

Known sets of gene markers associated with cardiovascular pathologies with high level of evidence (green marking) were retrieved from the PanelApp from Genomics England database (https://panelapp.genomicsengland.co.uk/) for gene sets enrichment analysis (GSEA) (panels: Arrhythmogenic right ventricular cardiomyopathy (Version 3.11), Brugada syndrome and cardiac sodium channel disease (Version 3.10), Cardiac arrhythmias (Version 13.37), Catecholaminergic polymorphic VT (Version 4.6), Dilated and arrhythmogenic cardiomyopathy (Version 2.31), Dilated Cardiomyopathy and conduction defects (Version 1.94), Familial non syndromic congenital heart disease (Version 1.86), Hypertrophic cardiomyopathy (Version 4.13), Idiopathic ventricular fibrillation (Version 1.2), Left Ventricular Noncompaction Cardiomyopathy (Version 1.4), Long QT syndrome (Version 3.8), Paediatric or syndromic cardiomyopathy (Version 4.10), Primary lymphoedema (Version 3.11), Progressive cardiac conduction disease (Version 2.8), RASopathies (Version 1.81), Sudden death in young people (Version 1.15), Thoracic aortic aneurysm or dissection (GMS) (Version 3.16)). Each set of genes was used to compute its respective enrichment in each cell of the dataset using AddModuleScore() from Seurat, scoring the average expression of the gene set compared to control sets of genes. The mean of the enrichment scores was computed per cluster for the coarse– and fine-grained annotation layers. The resulting mean values were visualized on heatmaps representing the gene set of interest across every cluster using geom_tile() from the ggplot2 package.

### Calculation of Single Cell Cluster Proportions

The relative sizes of coarse-grained clusters, as well as the fine-grained cluster representing FAPs, were assessed based on the ratio of cells in the cluster of interest, compared to the total number of cells in the relevant subset, and visualized with bar graphs in Microsoft Excel. The total number of cells in coarse– and fine-grained single cell clusters is displayed in Extended Table 10-11.

### Data Availability

All the data required to replicate the analysis, including cellranger output, spaceranger output, metadata, processed ISS data, extended figures and tables, as well as main RDS objects will be available on the Mendeley DATA repositories part 1 and part 2 upon publication. The processed data of Visium, scRNA-seq and ISS will be available for browsing gene expression, clusterings, and other analysis results on a viewer. The sensitive raw sequencing files for single-cell and spatial transcriptomics will be made available in the Federated European Genome-Phenome Archive upon publication and shared upon request. Due to the file size limitation, the corresponding author can provide access to the raw *in situ* sequencing dataset upon reasonable request. A list of abbreviations used in the manuscript, along with their expansions, is provided in Extended Table 12.

### Code Availability

The code used in this study, together with the material and information required to reproduce the Docker and Conda environments, will be shared through GitHub upon publication. A permanent version of the code will be uploaded to Zenodo upon publication.

## Supporting information

Ext. Tab. 12

Ext. Tab. 8

Ext. Tab. 9

Ext. Tab. 11

Ext. Tab. 10

Ext. Tab. 6

Ext. Tab. 7

Ext. Tab. 5

Ext. Tab. 3

Ext. Tab. 2

Ext. Tab. 4

Ext. Tab. 1

Ext. Text

## Acknowledgements

We acknowledge the services of the Developmental Tissue Bank of Karolinska Institute for providing prenatal tissue for this investigation. We thank Mariia Minaeva for her support with several bioinformatics tools used in this study, Quan H. Nguyen, Prakrithi Pavithra and Onkar Mulay for scientific discussions, Mattias Karlén for his help with illustrations, and Kim Thrane, Sami Sarenpää and Áron Kerényi for their useful suggestions on improving the text and figures of the manuscript. Finally, we thank the donors of tissue specimens whose invaluable contributions made this study possible.

## Author Contributions

Conceptualization: E.L., R.M., Ž.A., I.A., J. L.

Methodology: Ž.A., E.B., S.M.S., N.S., S.S., J.N.H.

Software: R.M., J.F., L.L., C.A., P.C.

Validation: S.M.S., N.S., S.S., J.N.H.

Formal analysis: E.L., R.M., Ž. A., J.F., M.H., L.L., M.V., P.C.

Investigation: Ž.A., E.B., S.M.S., N.S., S.S., J.N.H.

Resources: X.L., E.Lu., S.L., M.N., E.S., J.L.

Data curation: E.L., R.M., Ž.A., J.F., L.L., S.M.S., N.S., M.V., E.B.

Writing – original draft: E.L., R.M., Ž.A., J.F., S.M.S., N.S., S.S., J.N.H., E.B.

Writing – review and editing: E.L., R.M., Ž.A., J.L.

Visualization: E.L., R.M., Ž.A.

Supervision: E.L., O.B., C.S., E.Lu., S.L., M.N., E.S., I.A., J.L.

Project administration: E.L., R.M., Ž.A., J.L.

Funding acquisition: E.Lu., S.L., M.N., E.S., J.L.

## Supplemental Data Information

**Suppl. Fig. 1.**
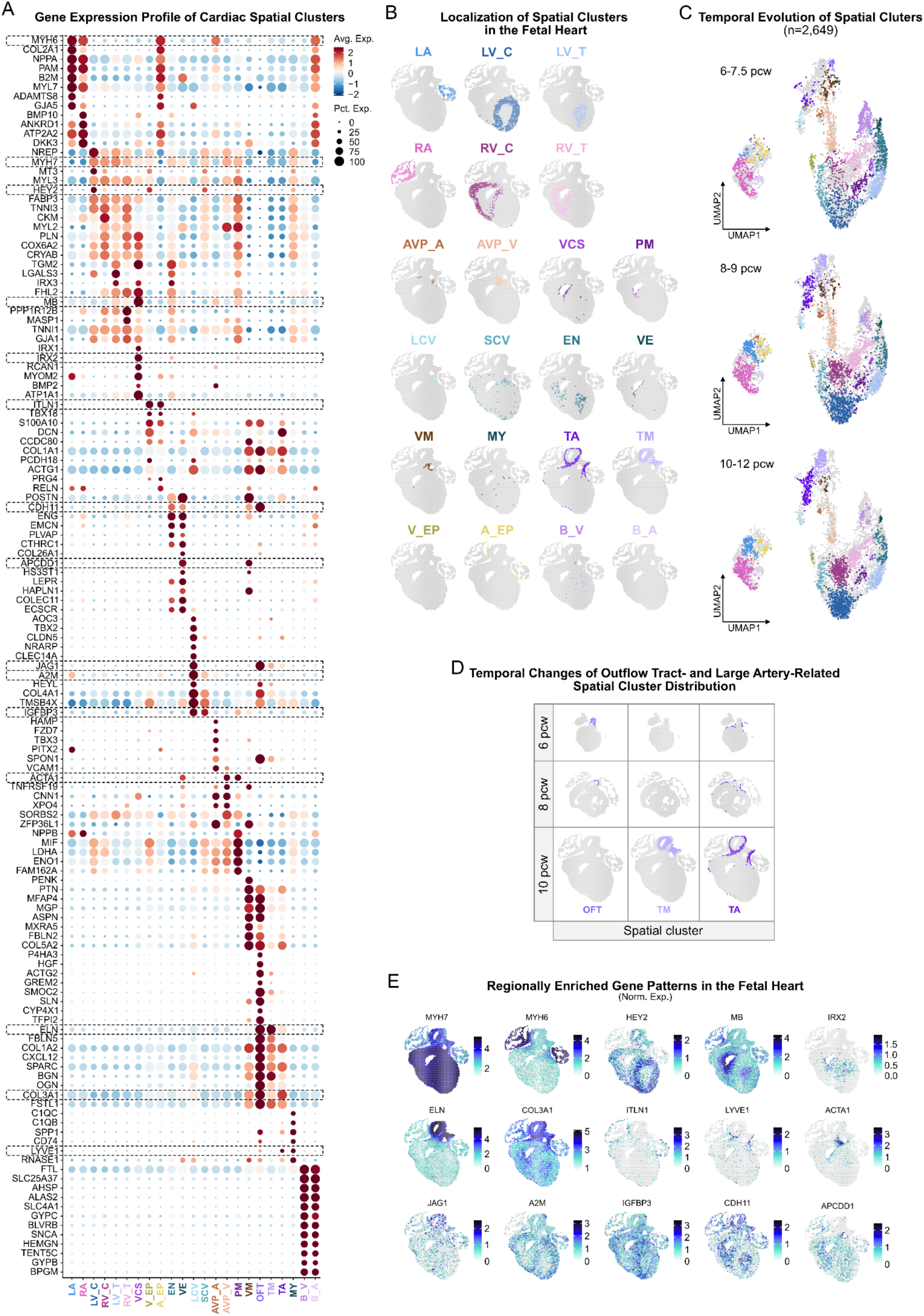
Transcriptomic Profiles and Temporal Transitions of Cardiac Spatial Clusters. **A**. *Gene Expression Profile of Cardiac Spatial Clusters.* Dot plot depicts the top 5 DEGs (log2FC > 0, p_val < 0.05) in the 23 spatial clusters. Genes highlighted by dashed rectangles were selected for spatial expression plots included in Suppl. Fig. 1e. **B.** *Localization of Spatial Clusters in the Fetal Heart.* The spatial cluster corresponding to the early outflow tract (OFT) is absent in the 10 pcw heart section. **C.** *Temporal Evolution of Spatial Clusters.* UMAPs illustrate the distribution of barcoded tissue spots (n=2,649) between spatial clusters across three developmental age groups (6-7.5, 8-9, 10-12 pcw). **D.** *Temporal Evolution of Outflow Tract– and Large Artery-Related Spatial Clusters.* The spatial cluster corresponding to the early outflow tract (OFT) gradually disappears, while the one representing the tunica media of large arteries (TM) expands by 10 pcw. **E.** *Regionally Enriched Gene Patterns in the Fetal Heart.* Spatial feature plots of selected DEGs (log2FC > 0, p_val < 0.05) of spatial clusters are presented in a 10 pcw heart section.

**Suppl. Fig. 2.**
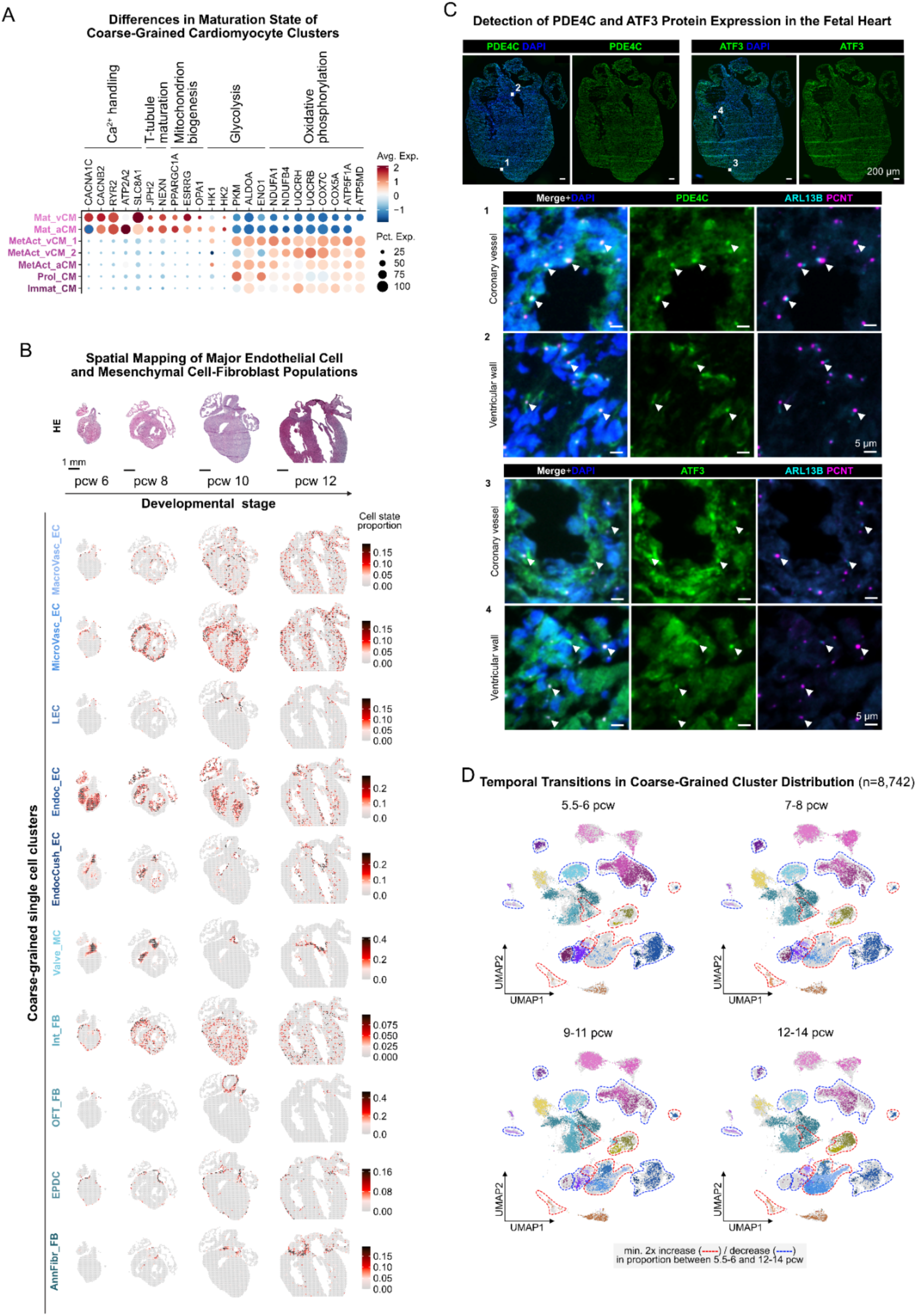
Molecular Characteristics and Temporal Transitions of Major Cardiac Cell Populations. **A**. *Differences in Maturation State of Coarse-Grained Cardiomyocyte Clusters.* Dot plot shows relative expression of genes involved in electromechanical activation and bioenergetics of cardiomyocytes in coarse-grained cardiomyocyte clusters. **B.** *Spatial Mapping of Major Endothelial Cell and Mesenchymal Cell-Fibroblast Populations.* The upper panel shows hematoxylin-eosin (HE) micrographs of 6, 8, 10 and 12 pcw heart sections, while the lower panels depict predicted cell state proportions of selected coarse-grained clusters in the corresponding Visium tissue spots. Scale bar represents 1 mm. **C.** *Detection of PDE4C and ATF3 Protein Expression in the Fetal Heart.* Immunofluorescence demonstrates widespread expression of PDE4C (upper left, green) and ATF3 (upper right, green) proteins in a 9 pcw heart section, along with DAPI nuclear staining (blue). Scale bar represents 200 μm. The middle and bottom panels feature ROIs including a coronary artery (1, 3) and part of the ventricular myocardium (2, 4), highlighting subcellular enrichment of PDE4C (middle, green) and ATF3 (bottom, green) proteins in basal bodies, along with PCNT (basal bodies, purple), ARL13B (primary cilia, cyan), and DAPI (nuclei, blue) staining. Scale bar represents 5 µm. Arrowheads indicate PDE4C/ATF3^+^-PCNT^+^ basal body structures with connected ARL13B^+^ primary cilia. **D.** *Temporal Transitions of Coarse-Grained Cluster Distribution.* UMAPs illustrate size changes of coarse-grained clusters across four developmental age groups (5.5-6, 7-8, 9-11, and 12-14 pcw, n=8,742). Dashed lines mark clusters with min. 2x increase (red) or decrease (blue) in proportion compared to the total number of cells between the 5.5-6 and 12-14 pcw age groups.

**Suppl. Fig. 3.**
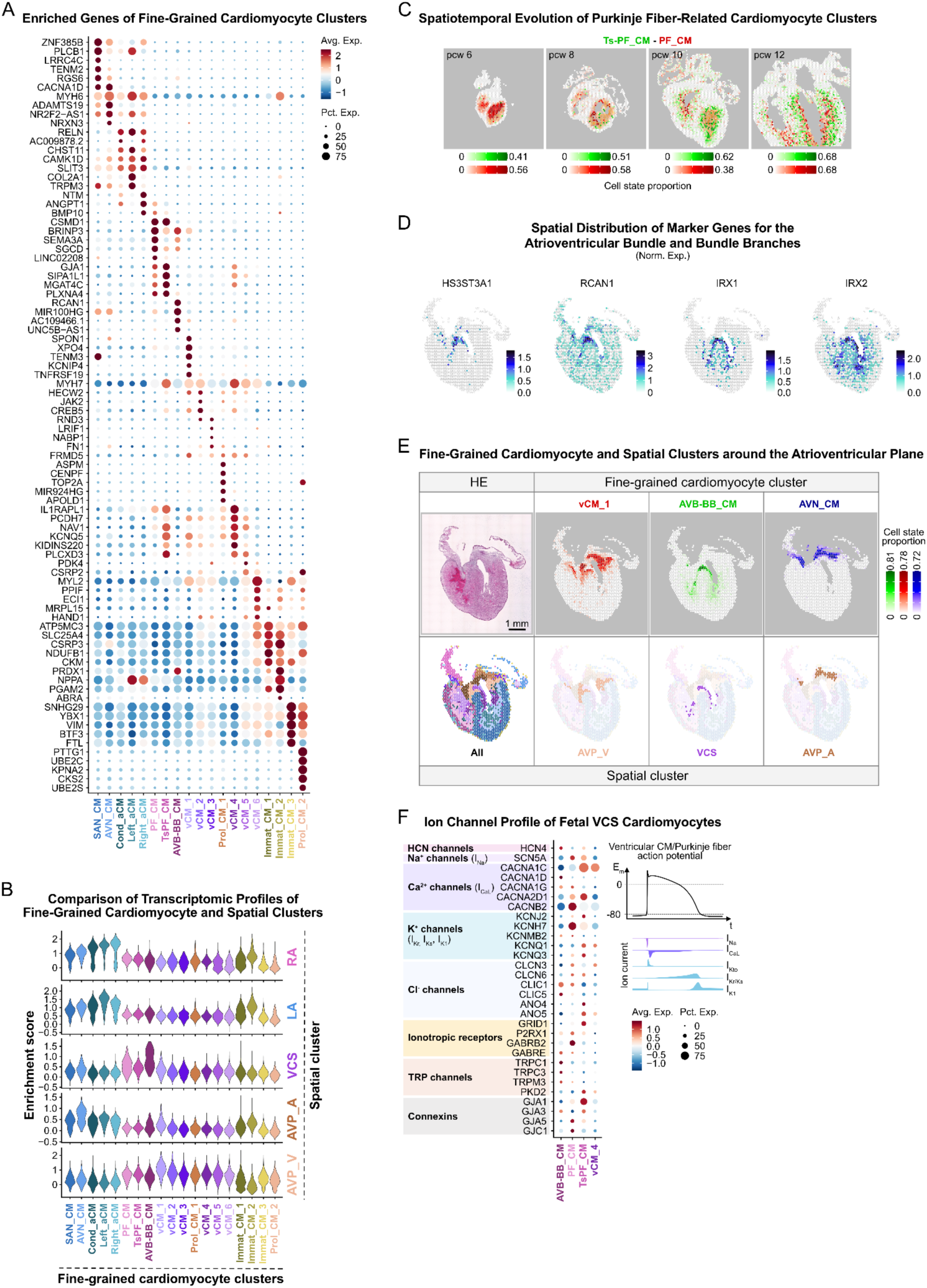
Molecular and Spatial Features of Fine-Grained Cardiomyocyte States. **A**. *Enriched Genes of Fine-Grained Cardiomyocyte Clusters.* Dot plot depicts the top 5 DEGs (log2FC > 0, p_val < 0.05) between the 19 clusters. **B.** *Comparison of Transcriptomic Profiles of Fine-Grained Cardiomyocyte and Spatial Clusters.* Violin plots show gene signature enrichment of selected spatial clusters in fine-grained cardiomyocyte cell states. Spatial clusters: LA–left atrial myocardium; RA–right atrial myocardium; VCS–ventricular conduction system; AVP_A–atrioventricular plane, atrial side; AVP_V– atrioventricular plane, ventricular side. **C.** *Spatiotemporal Evolution of Purkinje Fiber-Related Cardiomyocyte Clusters*. Predicted spatial distribution of Ts-PF_CMs (green) and PF_CMs (red) in 6, 8, 10 and 12 pcw heart sections supports their gradual separation. **D.** *Spatial Distribution of Marker Genes for the Atrioventricular Bundle and Bundle Branches.* Spatial feature plots display selected DEGs (log2FC > 0, p_val < 0.05) of AVB-BB_CMs in a 10 pcw heart section. **E.** *Fine-Grained Cardiomyocyte and Spatial Clusters around the Atrioventricular Plane.* The upper panels display the hematoxylin-eosin (HE) micrograph of a 10 pcw heart section, along with the predicted spatial distribution of vCM_1 (red), AVB-BB_CM (green) and AVN_CM (blue) cells, while the lower panels illustrate the highly similar localization of the AVP_V, VCS and AVP_A spatial clusters, respectively. Scale bar represents 1 mm. **f.** *Ion Channel Profiles of Cardiomyocytes in the Fetal Ventricular Conduction System.* Dot plot displays the relative expression of selected ion channel genes across the AVB-BB_CM, PF_CM, Ts-PF_CM, and the contractile vCM_4 cell states. Major ion currents of the ventricular cardiomyocyte/Purkinje fiber action potential are illustrated in colors consistent with the ion channel plot.

**Suppl. Fig. 4.**
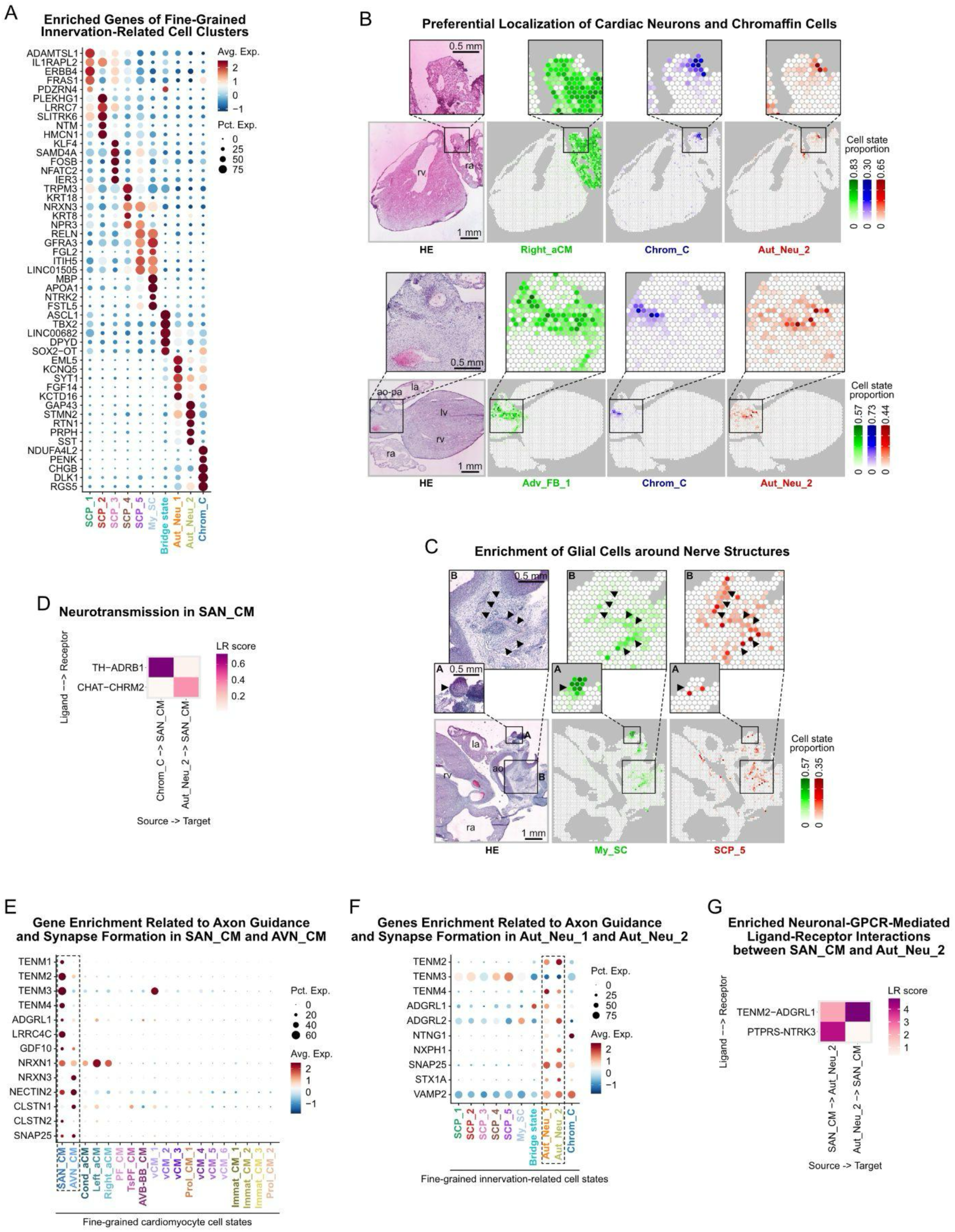
Transcriptomic and Spatial Characteristics of Fine-Grained Innervation-Related Cell States. **A**. *Enriched Genes of Fine-Grained Innervation-Related Cell Clusters.* Dot plot depicts the top 5 DEGs (log2FC > 0, p_val < 0.05) between the 10 clusters. **B.** *Preferential Localization of Neurons and Chromaffin Cells.* Chrom_C (blue) and Aut_Neu_2 cells (red) are closely associated with the atrial wall (outlined by Right_aCMs, green) and adventitia of the great arteries (outlined by Adv_FB_1 cells, green), presented in a 10.5 pcw (upper) and a 11 pcw (lower) heart section, along with their hematoxylin-eosin (HE) micrographs. ROIs highlight areas enriched in innervation-related cell states. **C.** *Enrichment of Glial Cells around Nerve Structures.* My_SCs (green) and SCP_5 cells (red) spatially associate with nerves (arrowheads) in the adventitia of the great arteries, highlighted in ROI A and B in the HE micrograph of a 11 pcw heart section. **D.** *Gene Expression Related to Axon Guidance and Synapse Formation across Fine-Grained Cardiomyocyte States.* Dot plot displays selective enrichment of relevant genes in SAN_CM and AVN_CM state. **E.** *Gene Expression Related to Axon Guidance and Synapse Formation across Fine-Grained Innervation-Related Cell States.* Dot plot displays selective enrichment of relevant genes in Aut_Neu_1 and Aut_Neu_2 cells. **F.** *Enriched Neural/GPCR-Mediated Ligand-Receptor Interactions between SAN_CM and Aut_Neu_2.* In panel D-E: la–left atrium, lv–left ventricle, ra–right atrium, rv–right ventricle, ao–aorta, pa–pulmonary artery; scale bars represent 1 mm in the main, and 0.5 mm in the zoom-in panels.

**Suppl. Fig. 5.**
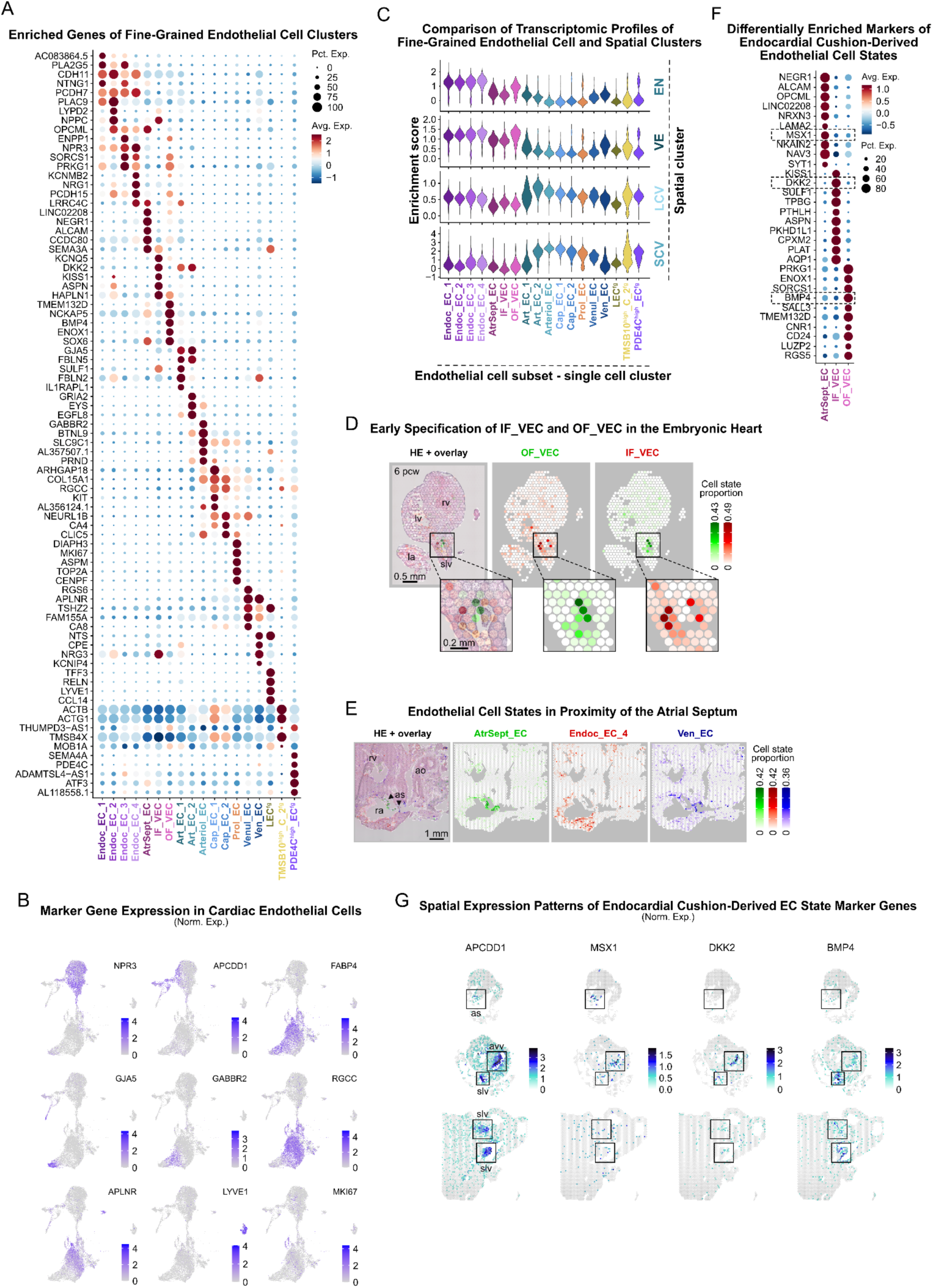
Molecular and Spatial Profiles of Fine-Grained Endothelial Cell States. **A**. *Enriched Genes of Fine-Grained Endothelial Cell Clusters.* Dot plot depicts the top 5 DEGs (log2FC > 0, p_val < 0.05) between the 18 clusters. **B.** *Marker Gene Expression in Cardiac Endothelial Cells.* Feature plots of selected DEGs (log2FC > 0, p_val < 0.05) outline endothelial cell populations of the endocardium (*NPR3*), endocardium-derived endothelial structures (*APCDD1*), great arteries and coronary vasculature (*FABP4*, *GJA5*, *GABBR2*, *RGCC*, *APLNR*), and lymphatic vessels (*LYVE1*), as well as proliferating cells (*MKI67*). **C.** *Comparison of Transcriptomic Profiles of Fine-Grained Endothelial Cell and Spatial Clusters.* Violin plots show gene signature enrichment of selected spatial clusters in fine-grained endothelial cell states. Spatial clusters: EN–endocardium and subendocardium; VE–valve endothelium; LCV–large coronary vessels; SCV–small coronary vessels. **D.** *Early Specification of IF_VEC and OF_VEC Cell States in the Embryonic Heart*. Spatial mapping traces IN_VECs (red) and OF_VECs (green) to opposite sides of the developing semilunar valves, identified in the hematoxylin-eosin (HE) micrograph of a 6 pcw heart section. Scale bars represent 0.5 mm in the main, and 0.2 mm in the zoom-in panels. **E.** *Endothelial Cell States in Proximity of the Atrial Septum.* Endoc_EC_4 cells (red) show spatial enrichment in the smooth-walled atrium, while Ven_ECs (blue) map to areas of venous character in the direct vicinity of the atrial septum outlined by AtrSept_ECs (green), visualized in a 11 pcw heart section. Scale bar on the HE micrograph represents 1 mm. **F.** *Differentially Enriched Markers of Endocardial Cushion-Derived Endothelial Cell States.* Dot plot displays the top 10 DEGs (log2FC > 0, p_val < 0.05) between AtrSept_EC, IF_VEC and OF_VEC clusters. Genes highlighted by dashed rectangles were selected for spatial expression plots included in Suppl. Fig. 5g. **G.** *Spatial Expression Patterns of Endocardial Cushion-Derived Endothelial Cell State Marker Genes.* Spatial feature plots visualize the expression of selected AtrSept_EC (*MSX1*), IF_VEC (*DKK2*) and OF_VEC (*BMP4*) marker genes in the atrial septum, atrioventricular and semilunar valves (marked by rectangles) in 8 (top, middle) and 12 pcw (bottom) heart sections. In panels D, E, and G: slv–semilunar valve, avv–atrioventricular valve, as–atrial septum, rv– right ventricle, lv–left ventricle, ra–right atrium, la–left atrium, ao–aorta.

**Suppl. Fig. 6.**
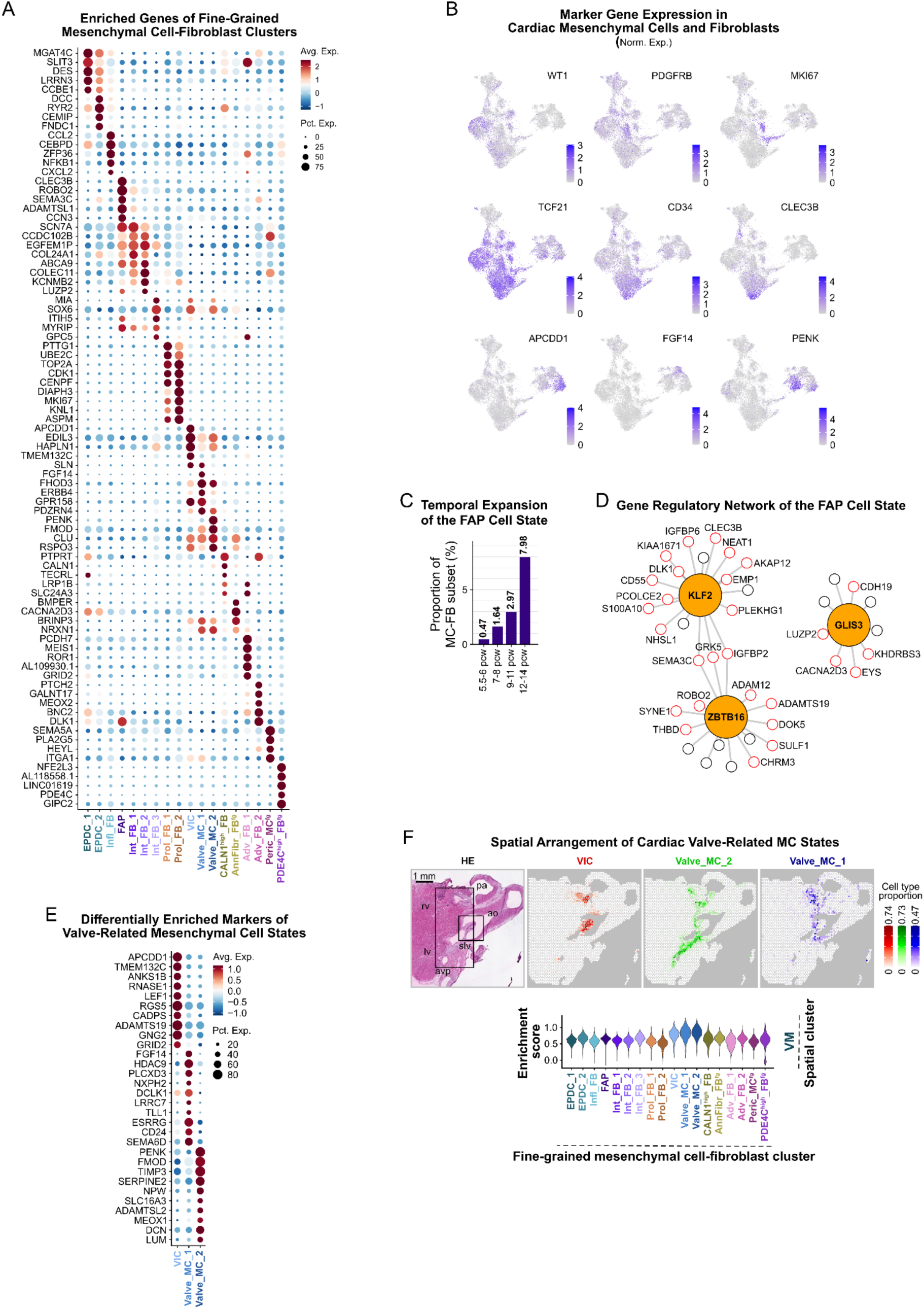
Transcriptomic and Spatial Features of Fine-Grained Fibroblast and Mesenchymal Cell States. **A**. *Enriched Genes of Fine-Grained Fibroblast and Mesenchymal Cell States.* Dot plot depicts the top 5 DEGs (log2FC > 0, p_val < 0.05) between the 18 clusters. **B.** *Marker Gene Expression in Cardiac Mesenchymal Cells and Fibroblasts.* Feature plots of selected DEGs (log2FC > 0, p_val < 0.05) outline cell populations with high expression of consensus epicardial (*WT1*), pericyte (*PDGFRB*), fibro-adipogenic progenitor (*CLEC3B*), putative cardiac fibroblast (*TCF21*, *CD34*), and proliferation (*MKI67*) markers, and genes enriched in distinct cardiac valve-related mesenchymal cell states (*APCDD1*, *FGF14*, *PENK*). **C.** *Temporal Expansion of the FAP Cell State.* Bar plot illustrates changes of the proportion of FAPs in the mesenchymal cell-fibroblast subset across four developmental age groups (5.5-6 pcw, 7-8 pcw, 9-11 pcw, 12-14 pcw). **D.** *Gene Regulatory Network of the FAP Cell State.* Differentially expressed transcription factors, presented with their associated target genes, in the FAPs. Orange-filled circles represent transcription factors, and red-rimmed circles represent DEGs (log2FC > 0, p_val < 0.05) compared to other fine-grained mesenchymal cell-fibroblast clusters. **E.** *Differentially Enriched Markers of Valve-Related Mesenchymal Cell States.* Dot plot displays the top10 DEGs (log2FC > 0, p_val < 0.05) between VIC, Valve_MC_1 and Valve_MC_2 clusters. **F.** *Spatial Arrangement of Cardiac Valve-Related Mesenchymal Cell States.* VICs (red) show spatial enrichment in the free segments of the sampled semilunar valves, while Valve_MC_1 (blue) and Valve_MC_2 cells (green) localize to the roots of the cusps. Valve_MC_2 also spreads out to larger segments of the intervalvular fibrous tissue towards the atrioventricular valves, while Valve_MC_1 is enriched around the semilunar valves (marked by rectangles) (upper). Scale bar on the hematoxylin-eosin (HE) micrograph represents 1 mm. Rv–right ventricle, lv–left ventricle, ao–aorta, pa–pulmonary artery, slv–semilunar valve, avp–atrioventricular plane. Violin plot shows enrichment of the valve mesenchyme (VM) spatial cluster gene signature in all three fine-grained mesenchymal cell clusters related to the cardiac valves (lower).

**Suppl. Fig. 7.**
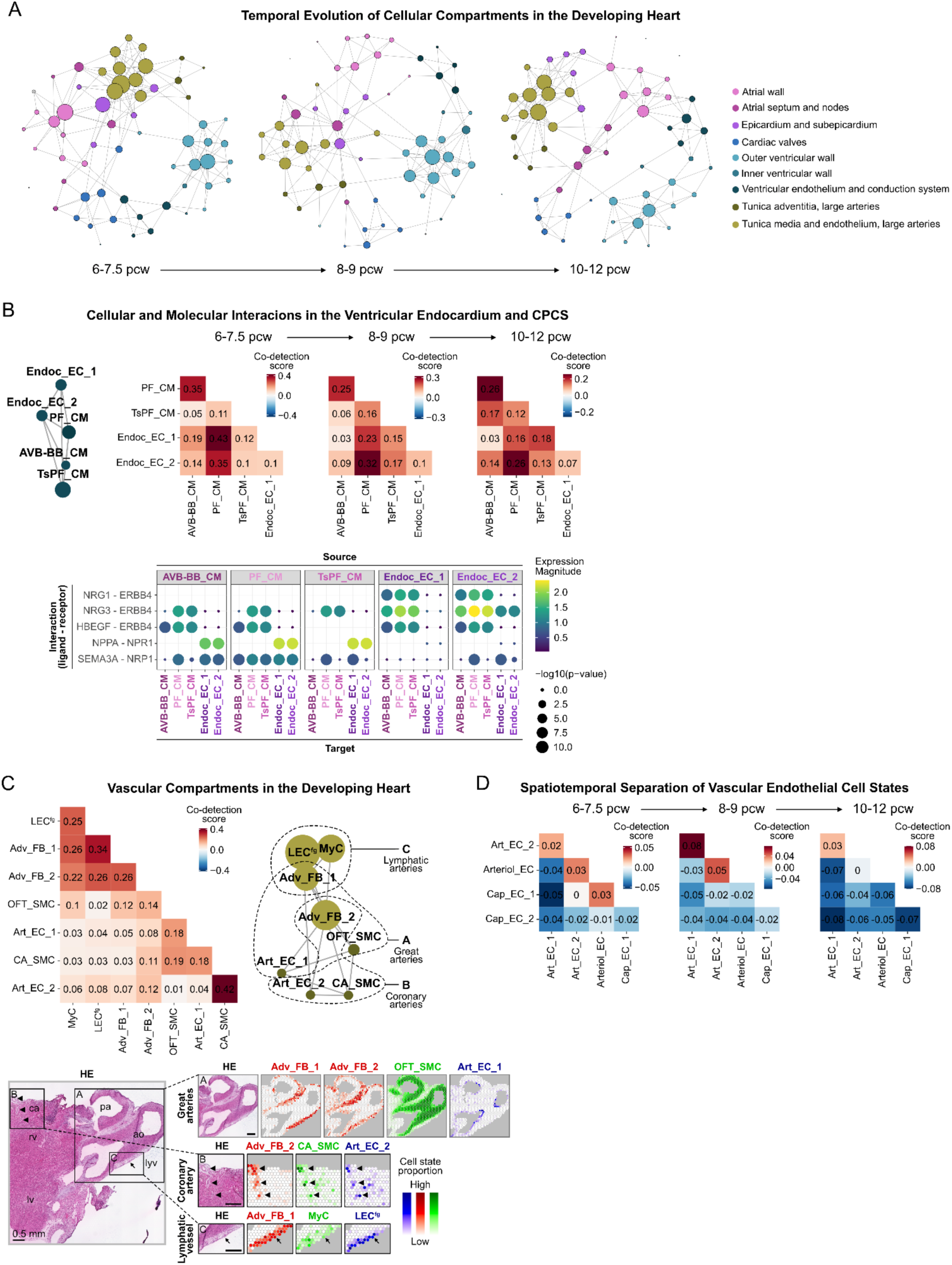
Temporal Transitions and Analysis of Selected Developmental Cardiac Niches. **A**. *Temporal Evolution of Cellular Compartments in the Developing Heart.* Niche network graphs calculated from cell state co-detection scores are displayed across three developmental age groups (6-7.5, 8-9, 10-12 pcw). Circles represent fine-grained cell states, and grey lines indicate their spatial association. Circle size reflects the number of closely associated cell states, and different colors indicate distinct cardiac structural compartments, consistently with Fig. 7A. **B.** *Cellular and Molecular Interactions in the Ventricular Endocardium and CPCS.* Niche network graph represents cellular components of the ventricular endocardium and conduction system (CPCS) (upper left), along with corresponding co-detection scores across three developmental age groups (6-7.5, 8-9, 10-12 pcw) (upper right). Selected ligand-receptor interactions, implicated in trabecular myocardium formation and Purkinje fiber specification, indicate functional differences between EndocEC_1 and Endoc_EC_2, and PF_CM and TsPF_CM, respectively (lower). **C.** *Vascular Compartments in the Developing Heart.* Niche network graph illustrates relations between cellular components of distinct cardiac vessel structures, outlined by dashed lines (upper right), with corresponding co-detection scores (upper left). Spatial mapping displays enrichment of Adv_FB_1 (red), Adv_FB_2 (red), OFT_SMC (green) and Art_EC_1 cells (blue) in the great arteries (ROI A), Adv_FB_2 (red), CA_SMC (green) and Art_EC_2 cells (blue) in coronary vessels (ROI B, marked by arrowheads), and Adv_FB_1 (red), MyC (green) and LEG^fg^ cells (blue) around lymphatic vessels (ROI C, marked by arrows). Scale bars represent 0.5 mm in both the main and zoom-in panels of hematoxylin-eosin (HE) micrographs. Rv–right ventricule, lv–left ventricule, ao–aorta, pa–pulmonary artery, ca– coronary artery, lyv–lymphatic vessel. **D.** *Spatiotemporal Separation of Vascular Endothelial Cell States.* Temporal changes of co-detection scores between vascular endothelial cell states are presented across three developmental age groups (6-7.5, 8-9, 10-12 pcw).

### Supplemental Table 1-7

Supplemental Table 1. List of Differentially Expressed Genes across Spatial Clusters

Supplemental Table 2. List of Differentially Expressed Genes across Coarse-Grained Single Cell Clusters

Supplemental Table 3. List of Differentially Expressed Genes across Fine-Grained Cardiomyocyte Clusters

Supplemental Table 4. List of Differentially Expressed Genes across Innervation-Related Cell Clusters

Supplemental Table 5. List of Differentially Expressed Genes across Fine-Grained Endothelial Cell Clusters

Supplemental Table 6. List of Differentially Expressed Genes across Fine-Grained Mesenchymal Cell-Fibroblast Clusters

Supplemental Table 7. Pearson Correlation-Based Co-Detection Scores for Fine-Grained Cell States

