## Supplementary material for "Spatial Dynamics of the Developing Human Heart": Ext. Text

**Extended Text - Additional Discussion**

**Sections:**

- Molecular Analysis of Mural Cell States
- Molecular Analysis of Blood-Related Cell States
- Transcriptomic Heterogeneity in Epicardium-Related Cell States
- Temporal Gene Expression Changes in Coarse-Grained Endothelial Cell and Mesenchymal Cell-Fibroblast Clusters
- Assessment of Spatiotemporal Transcriptomic Patterns in the Cardiac Valves, Outflow Tract and Great Arteries

### Molecular Analysis of Mural Cell States

Beyond the finely resolved and spatially characterized non-mural mesenchymal cell and fibroblast states, our single-cell dataset also includes coarse-grained clusters with smooth muscle and pericyte transcriptional characteristics (Fig. 2A, Ext. Fig. 2A). The two smooth muscle cell populations displayed distinct spatial patterns, one being enriched around the outflow tract and developing great vessels (OFT_SMC, 2938 cells), and the other found in a more extended localization within the myocardium with especially high proportion around the large coronary arteries (CA_SMC, 987 cells) (Fig. 2D, Suppl. Fig. 7C). Beyond a shared enrichment of genes consistent with smooth muscle identity (*MYH11*, *TAGLN*, *ACTA2*), the two populations featured vastly different transcriptomic profiles (Ext. Fig. 2B-C, Suppl. Table 2). As discussed in the 'Assessment of Spatiotemporal Transcriptomic Patterns in the Cardiac Valves, Outflow Tract and Great Arteries' section, OFT_SMCs shared several markers with the adventitial fibroblast population in the same region, OFT_FB, such as previously described regulators of outflow tract development (*MEIS1*[1], *PRDM6*[2,3], *LRP1B*[4]), but displayed selective enrichment of several extracellular matrix components (*ELN*, *FBLN5*) responsible for the elastic properties of the forming great arteries (Suppl. Table 2, Ext. Fig. 6A). These two gene groups displayed opposite temporal trends in our dataset, consistent with a shift from primary developmental processes towards maturation of the forming great arteries.

The CA_SMC cell state showed a large overlap in its transcriptomic profile, with even similar temporal trends, with the pericyte (PC, 2665 cells) population (Suppl. Table 2, Ext. Fig. 2B-C). This included classical markers of pericytes (*RGS5*, *KCNJ8*, *ABCC9*, *PDGFRB*, *CYGB*, *ENPEP*), displaying somewhat higher enrichment in the PC population. On the other hand, genes involved in local oxygen sensing (*HIGD1B*, *COX4I2*, *NDUFA4L2*) showed mildly stronger enrichment in the coronary smooth muscle cells, in line with these cells' role in mediating hypoxia-induced vasodilation in the coronary vasculature.[5] Pericytes have been previously proposed to serve as precursors for smooth muscle cells in the coronary vasculature, by upregulating *NOTCH3* at arterial remodeling sites in the vicinity of *JAG1*-expressing vascular endothelial cells, upon the onset of blood flow.[6] We observed consistent gene expression patterns in corresponding single-cell clusters with *NOTCH3* showing the highest enrichment in CA_SMC and PC, and *JAG1* in arterial endothelial cells (MacrVasc_EC) (Ext. Fig. 2D), and found high spatial enrichment of the *JAG1-NOTCH3* ligand-receptor pair around the large coronary and great arteries in our Visium dataset (Ext. Fig. 2E). These observations provide indirect support for a developmental connection between pericytes and coronary artery smooth muscle cells present in the human heart. Beyond shared markers, CA_SMCs also showed high expression of transcripts for the adhesion molecule catenin α3 (encoded by *CTNNA3*), the basal membrane component laminin α3 (encoded by *LAMA3*), the $Ca^{2+}$-binding protein  calsequestrin 2 (encoded by *CASQ2*), and gap junction protein-encoding *GJA4* (Ext. Fig. 2C).

Importantly, the pericyte population shared several highly enriched genes (*PLA2G5*, *SEMA5A*, *HEYL*, *ITGA11*, *GUCY1A1*, *CCDC3*, etc.) with a finely resolved cell state of the mesenchymal cell-fibroblast population, which we annotated as pericyte-like mesenchymal cells (Peric_MC$^{fg}$) (Ext. Fig. 2F). These cell states were mapped to largely complementary tissue regions, with PCs appearing in the compact ventricular myocardium, and Peric_MC$^{fg}$ cells in the atria and inner layers of the ventricular walls. Peric_MC$^{fg}$ had the highest expression of *APOE*, *ACTA2*, *IGFBP7*, and MCAM among the fine-grained mesenchymal cell-fibroblast cell states, consistent with a myofibroblast-like character (Suppl. Table 6). Cells with largely similar marker profiles have been annotated as myofibroblasts, as well as atrial fibroblasts in recent reports describing the cellular composition of the developing heart[7,8], highlighting the ambiguity regarding the precise identity of this population. Under healthy conditions, the adult human heart is largely devoid of myofibroblasts, which, however, readily differentiate from cardiac fibroblasts upon injury[9], thus the origin and role of cells with similar characteristics in the developing cardiac architecture is yet unclear.

**Molecular Analysis of Blood-Related Cell States**

In our single-cell dataset, we found a substantial number of blood-related cells, including red blood cells, immature platelets, and immune cells (Ext. Fig. 3A). Since red blood cells and platelets are not considered resident cellular components of the cardiac architecture, and appeared to be present in widely varying amounts between experimental samples, we decided to exclude them from downstream analysis, with the exception of cell state deconvolution of the Visium dataset, where the transcriptomic signature of these cell types still needs to be accounted for to obtain reliable results. We identified two coarse-grained clusters (marked as HL_excl_1 and HL_excl_4) dominated by red blood cell-specific transcripts (Suppl. Table 2, Ext. Fig. 3A-B). One of these clusters (HL_excl_1, 515 cells) showed stronger enrichment of embryonic hemoglobin isoforms (*HBE1*, *HBQ1*, *HBZ*, *HBM*), consistent with an immature embryonic/fetal erythrocyte phenotype, with the other one (HL_excl_4, 276 cells) being characterized by higher expression of components of the mature, adult-like hemoglobin complex (*HBA1*, *HBA2*, *HBB*). Furthermore, HL_excl_1 also included a small subset of cells expressing thrombocyte markers (*PF4*, *GP9*, *PPBP*, I*TGA2B*), presumably representing large embryonic platelets[10], and a small number of cells with granulocyte transcriptional characteristics (*IL5RA*, *FCER1A*) (Suppl. Table 2, Ext. Fig. 3B).

Myeloid and lymphoid cells were also present in our single-cell dataset (1593 and 667 cells, respectively) (Fig. 2A, Ext. Fig. 3A). The majority of myeloid cells featured markers consistent with yolk sac origin, such as *CX3CR1* and *CSF1R*, while a smaller population expressed *FLT3*, highlighting the gradual appearance of immune cells originating from the fetal liver[11] (Ext. Fig. 3C, upper panels). The distribution of these markers also aligned with the positions of $CCR2^-$ and $CCR2^+$ subpopulations within the myeloid compartment, respectively (data available in our interactive viewer). These results are consistent with observations in the adult heart, where *CCR2* expression was proposed to separate a macrophage population derived from definitive hematopoietic progenitors, constantly being renewed by monocyte recruitment, from self-renewing, tissue-resident macrophage population derived from the yolk sac under the fetal period.[12] The lymphoid population also appeared to be a mixture of cells with B-, T-, and NK-cell characteristics based on the expression pattern of *JCHAIN*, *CD3E* and *KLRB1* within the population (Ext. Fig. 3C, lower panels). While both the myeloid and lymphoid populations showed dispersed patterns throughout the tissue, co-detection analysis highlighted their highest spatial overlap with cells in the epicardial and subepicardial regions and the adventitia of the great vessels (Ext. Fig. 3D). These results align with the previously proposed seeding path for yolk sac-derived immune cells in mice, first populating a niche within and under the epicardial layer.[13] In addition, they provide indirect support to reports describing the central role of myeloid cells in the formation of coronary arteries and lymphatic vessels in the developing heart[14,15], by placing this cell type in the vicinity of such vessel structures (Suppl. Fig. 7C).

While we did not observe any distinct single-cell state with characteristics of lymphoid stromal cells, we captured high spatial enrichment of *CCL19*, a consensus marker of these cells, in several Visium sections, in concise segments within the subepicardium and adventitia of the great vessels (Ext. Fig. 3E). These areas showed a markedly high predicted proportion of lymphoid cells (LyC), and a close spatial association with lymphatic endothelial cells (LEC), supporting that the related tissue structures are, in fact, lymph nodes in the developing cardiac lymphatic system.

**Transcriptomic Heterogeneity in Epicardium-Related Cell States**

Several studies have described molecular heterogeneity within the developing epicardial compartment. However, human data related to this question are still scarce and partly derived from the analysis of iPSC-derived epicardioids[16–18].

In the developing heart, the epicardium serves as the source of a wide range of mesenchymal cells, including cardiac fibroblasts, pericytes and smooth muscle cells, while their potential contribution to the endothelial and cardiomyocyte compartments, while proposed by several animal studies under development and in tissue repair[19–23], is still largely debated. To generate mesenchymal cells, epicardial cells undergo epithelial-mesenchymal transition (EMT), during which they lose their epithelial characteristics, migrate into the subepicardial layer, and become multipotent intermediates called epicardium-derived progenitor cells (EPDCs), which subsequently give rise to the previously mentioned mesenchymal cell types. Many components of the EMT machinery are known, but because of the gradual nature of this process, there is no clear-cut molecular signature that defines the limit between epicardial and EPDC identities.

Epicardial heterogeneity discussed by previous human studies[16,17] was broadly related to the extent and activity of the EMT process in the analyzed epicardium-related samples. A recent report[16], comparing developmental and adult epicardial cells in single-cell datasets of dissociated whole heart tissue, identified three subpopulations within their epicardial compartment, including a mesothelial, a fibroblast-like, and a proliferating subpopulation, and highlighted temporal shifts, with the two latter populations disappearing in adult samples. Another work analyzed isolated human fetal subepicardial samples and identified two epithelial and three mesenchymal populations, however, molecular heterogeneity between clusters within the two groups was limited. These findings suggest that epicardial and epicardium-derived progenitor cells (at least in the earliest stages of their transition) were co-analyzed in these investigations. Taking this into consideration, it is plausible that inconsistencies between studies arise from shifts of cells representing the earliest stages of the EMT process between the epicardial and EPDC clusters. This is further supported by the fact that none of these studies define clear marker sets for the observed subpopulations, and rather describe them based on the relative and gradual enrichment of genes related to epicardial vs. mesenchymal identities. Furthermore, collection and processing of the analyzed heart samples might also differ (such as entire hearts vs. enriched epicardial and subepicardial tissue; processing with or without cell sorting; etc.), which might also contribute to the overall variability within the individual dataset, and thus to the definition of different clusters. Finally, it is challenging to differentiate between cellular heterogeneity inherently present in the resting epicardium and that due to a commenced differentiation process.

In agreement with these findings, we also observed an epicardial (EpC, 1187 cells) and a separate epicardium-derived progenitor cell (EPDC, 3837 cells) coarse-grained cluster in our dataset (Fig. 2A, Ext. Fig. 4A, Suppl. Table 2). Subsequent subclustering of non-mural mesenchymal cell-fibroblast subset allowed us to further refine cells with EPDC transcriptional characteristics, outlining two distinct fine-grained cell states, EPDC_1 and EPDC_2 (Fig. 6A, Suppl. Fig. 6A, Suppl. Table 6). While EpCs showed gene enrichment consistent with a mesothelial cell identity (*ITLN1*, *SBSPON*, *EZR*, *UPK3B*, *TNNT1*, *KRT19*, *BNC1*, *MSLN*, *PRG4*), EDPC_1 and EPDC_2 cell states also expressed mediators of the EMT process (*SPARC*, *POSTN*, *SNAI2*) and other relevant transcripts associated with epicardium-derived mesenchymal identity highlighted by other studies[8,16,17] (*MMP11*, *MOXD1*, *CCBE1*, *VEGFC*, *FNDC1*, etc.), beyond shared enrichment of the classically used epicardial markers *WT1* and *TBX18* (Ext. Fig. 4B-C). Notably, cell state mapping not only supported our annotations but also revealed spatial variations within the subepicardium between the two EPDC cell states. We found EPDC_1 being concentrated at the atrioventricular groove and EPDC_2 around the ventricular surface, potentially indicating regional differences in the mesenchymal transition process, while EpC appeared in its expected position, on the outer lining of the heart (Fig. 6C). Furthermore, while some epicardial markers exhibited relatively consistent expression throughout the investigated time frame in the EpC state, we observed a gradual temporal increase in others (*MSLN*, *C3*, *PRG4*), suggesting that these markers play a defining role in mature mesothelial characteristics (Ext. Fig. 4D).

Assessing the presence of a potentially epicardium-derived cardiomyocyte subpopulation in the developing heart is not feasible from the analyzed single-cell transcriptomics datasets, due to ambiguity regarding the relevant age range, overall low number of epicardial cells, and the high level of heterogeneity within the cardiomyocyte compartment. Importantly, cardiomyocyte-related gene expression in epicardial subsets[22] is not conclusive of actual cardiomyogenic differentiation, and previously published fate-mapping studies have been broadly challenged based on presumed leakiness of selected targeting approaches (such as WT1- or TBX18-based reporter gene expression[19–21], or pericardial injection of TAT-Cre recombinase[23]). Thus, and similarly to the other mentioned human studies, we could not observe any clear sign of a potential epicardial contribution to the cardiomyocyte compartment in our dataset.

**Temporal Gene Expression Changes in Coarse-Grained Endothelial Cell and Mesenchymal Cell-Fibroblast Clusters**

Temporally resolved differential gene expression analysis across coarse-grained single-cell clusters provided insight into relevant molecular transitions in the endothelial (Ext. Fig. 5A-B) and mesenchymal cell-fibroblast (Ext. Fig. 5A, C) populations, beyond highly enriched markers calculated from the integrated dataset (Suppl. Table 2).

Between the macro- (MacroVasc_EC) and microvascular endothelial cell clusters (MicroVasc_EC) of the developing coronary vasculature, we observed an opposite temporal pattern for *CLDN5*, *PRND*, and *IGFBP3* gene expression, implicated in the regulation of endothelial barrier function and angiogenesis, suggesting diverging characteristics related to these cellular functions from an early developmental stage. These populations displayed similar gradual enrichment of many shared marker genes over time, beyond some more specifically distributed and gradually enriched transcripts (*GJA5*, *GJA4*, *EYS*, *DKK2* in the MacroVasc_EC, and *CDH13*, *APLNR* in the MicroVasc_EC cluster). On the other hand, we detected strong selective enrichment of several marker genes of endocardial (Endoc_EC) (*PCDH7*, *TMEM100*, *CLEC3B*) and endocardial cushion-related endothelial cell clusters (EndocCush_EC) (*APCDD1*, *LTC4S*) already in the earliest analyzed age group (5.5-6 pcw), underscoring an even earlier specification between these populations (Ext. Fig. 5A-B).

In the mesenchymal cell-fibroblast subset, we focused our analysis on the interstitial fibroblast (Int_FB) and pericyte-like mesenchymal cell (Peric_MC) populations, which appeared to be the dominant mesenchymal cell components of the ventricular and atrial walls, respectively (consistent observation on the fine-grained level are presented in the niche network in Fig. 7A). Interestingly and in line with a recent report[7], we found the highest enrichment of *TCF21*, a previously proposed marker of epicardial and epicardium-derived progenitor cells, in the Int_FB population across the entire investigated time frame, supporting the notion that this gene's expression is associated with a more differentiated fibroblast identity in the developing human heart. At the same time, several consensus markers of cardiac fibroblasts (*DCN*, *LUM*, *C7*, *ABCA9*) showed gradual increase in expression in this population, along with *ROBO2*, a highly enriched gene in the fine-grained fibro-adipogenic progenitor-like (FAP) cell state identified in our dataset. This molecule is a component of the SLIT-ROBO signaling pathway, which is increasingly recognized in the modulation of fibrotic responses in various organs[24–26] (Ext. Fig. 5A, C).

The Peric_MC population, on the other hand, showed marked, although gradually decreasing enrichment of *THY1*, another proposed cardiac fibroblast marker, compared to other coarse-grained clusters of the mesenchymal cell-fibroblast subset. Importantly, the expression of this gene appeared to be highest in the pericyte population throughout the investigated timeframe, adding to several transcriptomic similarities between the two cell states (also highlighted for the fine-grained, Peric_MC$^{fg}$ population in Ext. Fig. 2F). In terms of extracellular matrix components, the Int_FB and Peric_MC clusters followed largely similar temporal trends and expression levels (*COL6A3*, *COL6A6*, *COL21A1*, *OGN*). On the other hand, *DCN* and *LUM*, encoding the the small leucine-rich repeat proteoglycans decorin and lumican, showed higher abundance in the Int_FB, and *TNC*, encoding tenascin C, in the Peric_MC population, highlighting compositional differences between the developing atrial and ventricular interstitial extracellular matrix (Ext. Fig. 5A, C).

Temporally differentially expressed genes in the mesenchymal cell population located in the outer annulus fibrosus (AnnFibr_FB) showed substantial overlaps with other mesenchymal cell clusters, such as *BRINP3* and *MAGI2* (also gradually downregulated in Valve_MC cluster) or *FBLN1* (showing similar temporal enrichment in the EPDC and OFT_FB clusters) (Ext. Fig. 5A, C).

Temporal gene expression patterns related to the OFT_FB and Valve_MC coarse-grained clusters, as well as the OFT_SMC, CA_SMC and PC mural cell populations are presented in Extended Figure 6A-C and Extended Figure 2A-F and the related text, respectively.

**Assessment of Spatiotemporal Transcriptomic Patterns in the Cardiac Valves, Outflow Tract and Great Arteries**

The Visium dataset in our study provided comprehensive spatial gene expression information from various stages of early heart development, enabling the exploration of spatiotemporal expression patterns during cardiogenesis. However, analyzing gene expression independently within spatial clusters or regions is not advisable due to several technical challenges, such as differences in cell type composition and density per spot, and discrepancies between sampled anatomical regions. A more robust approach involves integrating the spatial component through spatially aware annotation of cell states identified in the independent single-cell RNA-sequencing dataset. By selecting the dominant cellular components of regions of interest, these can be assessed through time-resolved differential gene expression analysis to uncover relevant molecular transitions, which can then be spatially validated in the Visium dataset in sections representing consecutive developmental stages. This strategy leverages the overall rich single-cell RNA-sequencing data, providing a more detailed assessment than independent analysis of spatial datasets would allow.

Accordingly, we assessed temporal transcriptional changes in selected regions of interest by utilizing results of time-resolved differential gene expression analysis across age-subsetted (5.5-6 pcw, 7-8 pcw, 9-11 pcw, and 12-14 pcw) subpopulations of all coarse-grained single-cell clusters. We deduced spatiotemporal patterns related to the outflow tract and great arteries, as well as the cardiac valves, by investigating highly expressed, temporally differentially expressed genes identified in the OFT_FB, OFT_SMC, and Valve_MC single-cell clusters (Ext. Fig. 6A), which show highly specific localization in these regions (Fig. 2D, Suppl. Fig. 2B). In the OFT_FB and OFT_SMC populations, we found several previously proposed mediators of outflow tract development related to the neural crest origin of certain cellular components of this region, such as *PRDM6*[2,3], *MEIS1*[1], *LRP1B*[4], showing decreasing expression over time. Simultaneously, key determinants of the developing great vessels' elastic properties (*ELN*, *PI15*) became gradually enriched in the OFT_SMC cluster throughout the investigated timeframe, with a similar trend in non-canonical NOTCH ligand *DLK1* in the OFT_FB cluster, presumably representing epicardium-derived contribution to this population.[27] In the Valve_MC cluster, we observed a clear temporal downregulation of *SEMA3D* and an upregulation of *COL12A1*, suggesting distinct roles for these molecules in the early formation and subsequent maturation of cardiac valves.[28,29]

Importantly, we spatially validated the observed expression patterns in sections included in our Visium dataset (Ext. Fig. 6B-C), confirming the reliability of our approach to deduce spatiotemporal molecular signatures from the temporal analysis of spatially annotated cell states. This strategy can be refined by applying the same temporal differential gene expression analysis on finely resolved single-cell clusters with distinct spatial distributions, as outlined by the predicted cellular composition of cardiac compartments and niches (Fig. 7A). However, low cell numbers in certain time-resolved clusters can be a limiting factor in this type of analysis.
